# Convergent Evolution of Conserved Mitochondrial Pathways Underlies Repeated Adaptation to Extreme Environments

**DOI:** 10.1101/2020.02.24.959916

**Authors:** Ryan Greenway, Nick Barts, Chathurika Henpita, Anthony P. Brown, Lenin Arias Rodriguez, Carlos M. Rodríguez Peña, Sabine Arndt, Gigi Y. Lau, Michael P. Murphy, Lei Wu, Dingbo Lin, Jennifer H. Shaw, Joanna L. Kelley, Michael Tobler

**Affiliations:** Division of Biology, Kansas State University, Manhattan, KS, USA; Department of Integrative Biology, Oklahoma State University, Stillwater, OK, USA; School of Biological Sciences, Washington State University, Pullman, WA, USA; División Académica de Ciencias Biológicas, Universidad Juárez Autónoma de Tabasco, Villahermosa, Tabasco, Mexico; Instituto de Investigaciones Botánicas y Zoológicas, Universidad Autónoma de Santo Domingo, Santo Domingo, Dominican Republic; MRC Mitochondrial Biology Unit, University of Cambridge, Cambridge, United Kingdom; Department of Biosciences, University of Oslo, Oslo, Norway; Department of Nutritional Sciences, Oklahoma State University, Stillwater, OK, USA; Department of Biomedical Sciences, PCOM South Georgia, Moultrie, GA, USA

**Keywords:** Adaptive evolution, Comparative physiology, Ecological genomics, Hydrogen sulfide, Phylogenetic comparative analysis, Poeciliidae

## Abstract

Extreme environments test the limits of life; yet, some organisms thrive in harsh conditions. Extremophile lineages inspire questions about how organisms can tolerate physiochemical stressors and whether the repeated colonization of extreme environments is facilitated by predictable and repeatable evolutionary innovations. We identified the mechanistic basis underlying convergent evolution of tolerance to hydrogen sulfide (H_2_S)—a toxicant that impairs mitochondrial function—across evolutionarily independent lineages of a fish (*Poecilia mexicana*, Poeciliidae) from H_2_S-rich springs. Using comparative biochemical and physiological analyses, we found that mitochondrial function is maintained in the presence of H_2_S in sulfide spring *P. mexicana*, but not ancestral lineages from nonsulfidic habitats, due to convergent adaptations in the primary toxicity target and a major detoxification enzyme. Genome-wide local ancestry analyses indicated that convergent evolution of increased H_2_S tolerance in different populations is likely caused by a combination of selection on standing genetic variation and *de novo* mutations. At a macroevolutionary scale, H_2_S tolerance in 10 independent lineages of sulfide spring fishes across multiple genera of Poeciliidae is correlated with the convergent modification and expression changes of genes associated with H_2_S toxicity and detoxification. Our results demonstrate that the modification of highly conserved physiological pathways associated with essential mitochondrial processes mediates tolerance to physiochemical stress. In addition, the same pathways, genes, and—in some instances—codons are implicated in H_2_S adaptation in lineages that span 40 million years of evolution.

**Significance Statement:** Some organisms can tolerate environments lethal for most others, but we often do not know what adaptations allow them to persist and whether the same mechanisms underly adaptation in different lineages exposed to the same stressors. Investigating fish inhabiting springs rich in toxic hydrogen sulfide (H_2_S), we show that tolerance is mediated by the modification of pathways that are inhibited by H_2_S and those that can detoxify it. Sulfide spring fishes across multiple genera have evolved similar modifications of toxicity targets and detoxification pathways, despite abundant lineage-specific variation. Our study highlights how constraints associated with the physiological consequences of a stressor limit the number of adaptive solutions and lead to repeatable evolutionary outcomes across organizational and evolutionary scales.

---

Stephen J. Gould made a strong case for the importance of contingency in evolution, famously quipping that replaying the “tape of life” would lead to different outcomes every time (1). But despite the unpredictability of mutations, the effects of genetic drift, and other historical contingencies, convergent evolution of phenotypic traits and their underlying genes is common, indicating that natural selection sometimes finds repeatable and predictable solutions to shared evolutionary challenges (2, 3). A major challenge that remains is the identification of the ecological, genetic, and functional factors that might determine the repeatability and predictability of evolutionary outcomes (4).

Mitochondria and their genomes provide a fascinating model to ask questions about the predictability of evolution for two reasons: (*i*) Mitochondrial genomes were historically thought to be a prime example of contingency evolution, because alternative genetic variants were assumed to be selectively neutral (5). This paradigm has been shifting though, with mounting evidence that mitochondria—and genes encoded in the mitochondrial genome—can play important roles in adaptation, especially in the context of physiochemical stress (6). (*ii*) Mitochondria are critical for the cellular function of eukaryotes (7). Their function is dependent on the gene products from two genomes, the mitochondrial and the nuclear (8), which interact to ultimately shape whole-organism performance. Despite extensive characterization of allelic variation in mitochondrial genomes, it often remains unclear how variation in genes that contribute to mitochondrial function translates to variation in physiological and organismal function. Furthermore, it is not known whether exposure to similar selective regimes may cause convergent modifications of mitochondrial genomes and emergent biochemical and physiological functions in evolutionarily independent lineages.

Extreme environments that represent novel ecological niches are natural experiments to address questions about mechanisms underlying mitochondrial adaptations and illuminate the predictability of adaptive evolution of mitochondria. Among the most extreme freshwater ecosystems are springs with high levels of hydrogen sulfide (H_2_S), a potent respiratory toxicant lethal to metazoans due to its inhibition of mitochondrial ATP production (9). Multiple lineages of livebearing fishes (Poeciliidae) have colonized H_2_S-rich springs throughout the Americas and independently evolved tolerance to sustained H_2_S concentrations orders of magnitude higher than those encountered by ancestral lineages in nonsulfidic habitats (10). Here, we identify the molecular basis of an evolutionary innovation that facilitated the independent colonization of extreme environments (increased H_2_S tolerance) and ask if the underlying mechanisms have evolved in convergence in disparate lineages of livebearing fishes.

H_2_S toxicity and detoxification are associated with highly conserved physiological pathways in mitochondria (Figure 1A) (11, 12), providing *a priori* predictions about the potential molecular mechanisms underlying adaptation to this strong source of selection. Toxic effects of H_2_S result from binding to and inhibition of cytochrome c oxidase (COX) in the oxidative phosphorylation (OxPhos) pathway, which contains subunits encoded in both the nuclear and the mitochondrial genomes (13). Animal cells can also detoxify low concentrations of endogenously produced H_2_S via the mitochondrial sulfide:quinone oxidoreductase (SQR) pathway, which is linked to OxPhos but entirely encoded in the nuclear genome (14). We have previously shown that genes associated with both pathways are under divergent selection and differentially expressed between fish populations in sulfidic and nonsulfidic habitats (10). These include nuclear and mitochondrial genes encoding subunits of the direct toxicity target (COX) and the nuclear gene encoding the enzyme mediating the first step of detoxification (SQR) (10). Tolerance to H_2_S may therefore be mediated by resistance (modification of toxicity targets that reduce the negative impact of H_2_S), regulation (modification of physiological pathways that maintain H_2_S homeostasis), or both (9).

**Figure 1.**
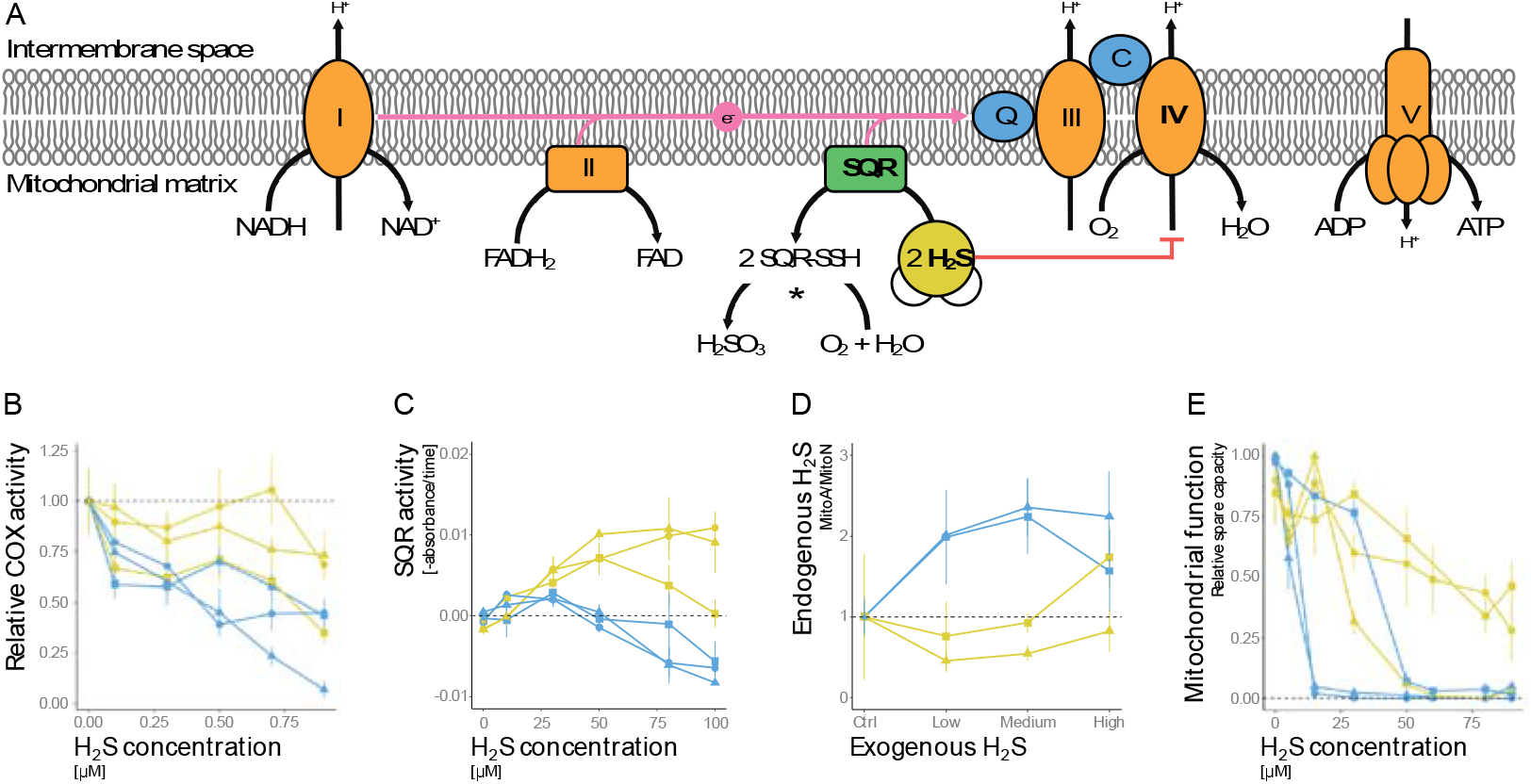
**A.** Physiological pathways associated with H_2_S toxicity and detoxification are located in the inner mitochondrial membrane. H_2_S inhibits OxPhos (orange enzymes, encoded by genes in the mitochondrial and nuclear genomes) by binding to COX (Complex IV). H_2_S can be detoxified through SQR (green enzyme, encoded by genes in the nuclear genome) and additional enzymes (indicated by asterisks). **B.** Relative activity of COX upon H_2_S exposure, which was primarily explained by an interaction between habitat type of origin and ambient H_2_S concentration (Tables S2-S3). **C.** Activity of SQR as a function of H_2_S concentration, which was explained by an interaction between habitat type of origin and H_2_S concentration (Tables S4-S5). **D.** Relative change in mitochondrial H_2_S concentrations in the liver of live fish exposed to different levels of environmental H_2_S. Variation in mitochondrial H_2_S levels were explained by habitat type of origin and exogenous H_2_S concentration (Tables S6-S7). **E.** Relative spare respiratory capacity of isolated liver mitochondria at different levels of H_2_S. The interaction between habitat type of origin and drainage of origin best explained variation in spare respiratory capacity (Tables S11-S12). For all graphs, yellow colors denote *P. mexicana* from H_2_S-rich habitats, blue from nonsulfidic habitats. Symbols stand for populations from different river drainages (■: Tac; ▲: Puy; ●: Pich; see Figure S1).

Based on these previous results, we hypothesized that the repeated modification of enzymes in the OxPhos and SQR pathways in *P. mexicana* populations from sulfidic habitats leads to an increased ability to maintain mitochondrial function in the presence of H_2_S. In the present study, we consequently used a series of *in vivo* and *in vitro* assays to identify the functional consequences of modifications to the OxPhos and SQR pathways in evolutionarily independent population pairs of *P. mexicana* from adjacent sulfidic and nonsulfidic habitats that are situated in different river drainages. In addition, we hypothesized that convergent molecular modifications in the same pathways underlie the convergent evolution of H_2_S tolerance across different lineages of poeciliid fishes. Hence, we also used phylogenetic comparative analyses of gene expression and analyses of molecular evolution to detect patterns of molecular convergence in 10 lineages of sulfide spring poeciliids and ancestors from nonsulfidic habitats.

## Results and Discussion

### Sulfide spring *P. mexicana* exhibit a resistant toxicity target

If resistance is the primary mechanism of tolerance, we would predict that COX function is maintained in the presence of H_2_S in fish from sulfidic populations, but not those from nonsulfidic populations. Quantification of COX function indicated that enzyme activity generally declined with increasing H_2_S concentrations, but this decline was reduced in populations from sulfidic habitats (Figure 1B; habitat × H_2_S: *P* < 0.001, Table S3). Even though the drainage of origin was not retained as an explanatory variable in statistical models (SI Appendix, Table S2), COX activity in one H_2_S-tolerant population (Tac) declined just as in nonsulfidic populations (Figure 1B). The other two *P. mexicana* populations from sulfidic habitats (Puy and Pich) maintained significant COX activity even at the highest H_2_S concentrations, which should reduce the negative impact of H_2_S on cellular respiration. These results are consistent with previous analyses (15) and indicate that resistance may contribute to H_2_S tolerance in some populations, but cannot explain the repeated evolution of H_2_S tolerance by itself.

### Sulfide spring *P. mexicana* can regulate mitochondrial H_2_S through increased detoxification

We also tested whether tolerant and intolerant populations differ in their ability to detoxify H_2_S by conducting enzyme activity assays of SQR. Activity of SQR was significantly higher in mitochondria from sulfidic populations at intermediate and high H_2_S concentrations (Figure 1C; habitat × H_2_S: *P* < 0.001 in SI Appendix, Table S5), likely helping fish from sulfidic habitats to maintain H_2_S homeostasis during environmental exposure. To test this prediction *in vivo*, we used a novel mitochondria-specific H_2_S-probe (MitoA) that allows for the monitoring of relative H_2_S levels inside the mitochondria of living organisms (16). We measured mitochondrial H_2_S concentrations in this manner using laboratory-reared fish that were exposed to varying levels of environmental H_2_S. Because laboratory-reared fish were not available for the population pair from Pich, only two population pairs were used for this analysis. Overall, mitochondrial H_2_S concentrations increased with environmental exposure (*P* = 0.001) and was higher in fish from non-sulfidic habitats (P < 0.001 in SI Appendix, Table S7). H_2_S concentrations in mitochondria isolated from livers (Figure 1D) and other organs (SI Appendix, Figure S2) of fish from nonsulfidic habitats increased above control levels at all exposure concentrations. In contrast, mitochondrial H_2_S concentrations in isolates of fish from sulfidic populations did not usually exceed control levels and remained lower than levels in fish from nonsulfidic habitats. Together, these results indicate that populations of *P. mexicana* from sulfidic habitats can detoxify H_2_S at higher rates and thus regulate mitochondrial H_2_S upon environmental exposure.

### Sulfide spring *P. mexicana* can maintain mitochondrial function in presence of H_2_S

Modification of the OxPhos and SQR pathways in *P. mexicana* suggests that mitochondrial adaptations are key to the evolution of H_2_S tolerance. Therefore, mitochondrial function of fish from sulfidic habitats should be maintained upon exposure to H_2_S. We tested this hypothesis by quantifying different aspects of mitochondrial function (basal respiration, maximal respiration, and spare respiratory capacity) along a gradient of H_2_S concentrations using an *ex vivo* coupling assay. As expected, all aspects of mitochondrial function generally declined with increasing H_2_S (Figures 1E; SI Appendix, S3-S5). Comparison of mitochondrial function between adjacent populations in sulfidic and nonsulfidic habitats indicated no differences in basal respiration (SI Appendix, Figure S3). However, individuals from sulfidic populations were able to maintain maximal respiration and spare respiratory capacity at higher levels compared to individuals from nonsulfidic habitats of the same river drainage (Figure 1E), even though the magnitude of difference and the shape of response curves varied (SI Appendix; significant drainage × habitat interactions in Tables S10 & S12, Figures S4-S5). These findings indicate that mitochondria of H_2_S-tolerant individuals continue to produce ATP in the presence of a potent inhibitor that reduces mitochondrial function in ancestral lineages.

Overall, our quantitative analyses indicate clear patterns of convergence in functional physiological traits associated with H_2_S tolerance. Nonetheless, further inspection of the results also reveals lineage-specific patterns (especially in H_2_S-dependent COX activity and mitochondrial respiration), indicating that evolutionary responses across lineages are similar but not necessarily identical. These idiosyncrasies are consistent with the results of previous comparative transcriptome analyses, which revealed a large number of genes that are under selection or differentially expressed in just a subset of lineages in addition to genes that are consistently differentially expressed and under selection across all lineages (17–19). Based on their functions, the OxPhos and SQR pathways undoubtedly include some major-effect genes influencing H_2_S tolerance in different populations of *P. mexicana*, but tolerance—as an emergent physiological trait—is a complex trait impacted by other genes as well. In the future, quantitative genetic analyses will be required to understand how other loci contribute to tolerance within each lineage, and how population-specific patterns of genetic differentiation might shape variation in functional physiology evident in our data.

### Convergence among *P. mexicana* populations is shaped by selection on de novo mutations and standing genetic variation

The convergent evolution of H_2_S tolerance in *P. mexicana* begs questions about the origin of adaptive alleles (20). At microevolutionary scales, convergence may be a consequence of the repeated assembly of related alleles into different genomic backgrounds, either through selection on standing genetic variation or introgression (21, 22). However, the epitome of convergent evolution is, arguably, the independent origin of adaptive mutations at the same locus that lead to consistent functional outcomes (23). To identify convergence at a genomic level, we re-sequenced whole genomes of multiple *P. mexicana* individuals from sulfidic and nonsulfidic habitats. Analyzing phylogenetic relationship among *P. mexicana* populations (with *P. reticulata* as an outgroup) using 13,390,303 single nucleotide polymorphisms (SNPs) distributed across the genome confirmed three independent colonization events of sulfide springs and distinct evolutionary trajectories for sulfide spring populations in different drainages (Figure 2A), as inferred by previous studies (24). If adaptive alleles arose separately through *de novo* mutation in each sulfide spring population, we would expect that putative adaptive alleles mirror these relationships, as previously documented for H_2_S-resistant alleles in mitochondrial COX subunits (15). However, patterns of divergence (SI Appendix, Figure S6) and local ancestry were highly variable across the genome. Classifying local patterns of genetic similarity using a Hidden Markov Model and a self-organizing map allowed us to identify genomic regions in which ancestry patterns deviate from the genome-wide consensus, including multiple regions with a strong signal of clustering by ecotype (sulfidic *vs*. nonsulfidic populations). Such clustering by ecotype occurred in less than 1 % of the genome (SI Appendix, Figure S7), but included genomic regions encoding key genes associated with H_2_S detoxification (e.g., SQR and ETHE1; Figure 2B; SI Appendix, Dataset S2). Clustering by ecotype indicates a monophyletic origin of putatively adaptive alleles at these loci that are shared across independent lineages of sulfide spring *P. mexicana* as a consequence of selection on standing genetic variation or introgression (25), although the latter scenario is less likely considering the geographic barriers and strong survival selection against migrants from sulfidic to nonsulfidic habitats (26). Consequently, multiple mechanisms—not just selection on *de novo* mutations (19)—played a role in the convergent evolution of H_2_S-tolerance in *P. mexicana*.

**Figure 2.**
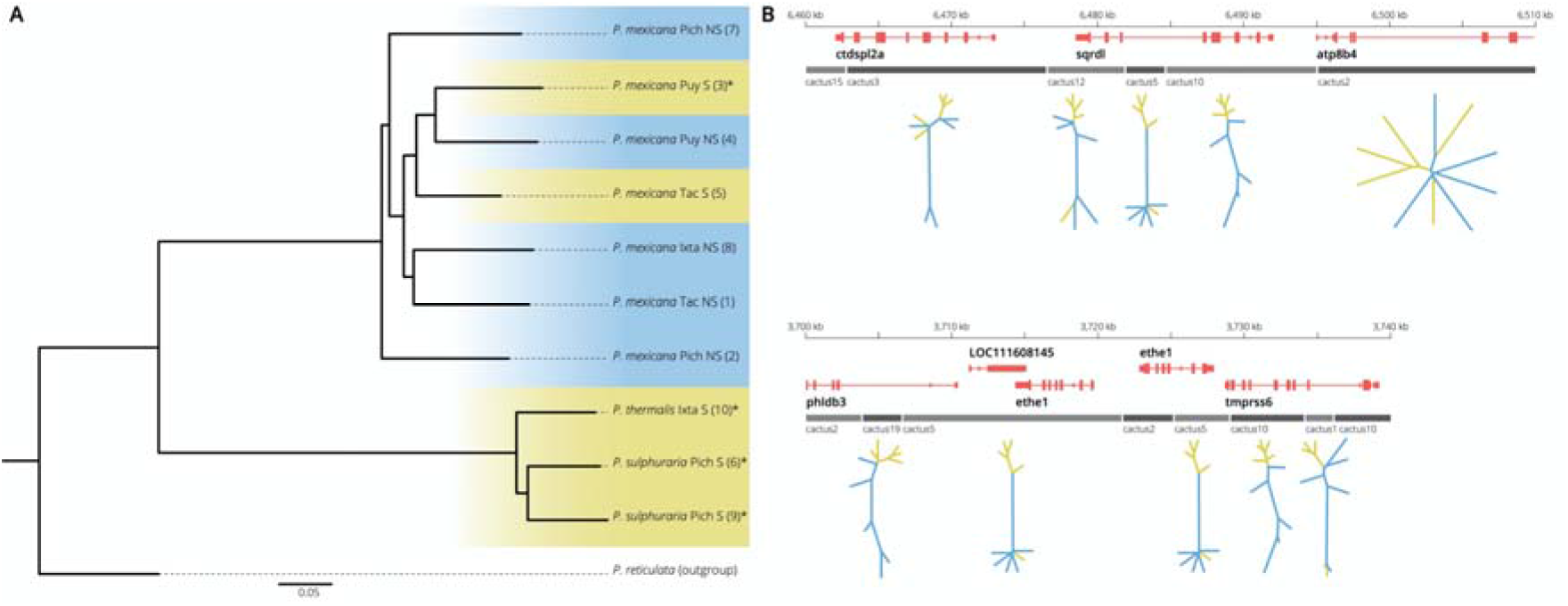
**A.** Phylogeny of different population in the *P. mexicana* species complex (with *P. reticulata* as an outgroup) based on genome-wide SNPs. Colors indicate sulfidic (yellow) and nonsulfidic (blue) lineages. **B.** Local ancestry patterns around genes encoding two enzymes involved in H_2_S detoxification, *SQR* and *ETHE1*. Gray bars represent the local ancestry pattern (cactus) associated with each region. Unrooted trees represent local ancestry relationships, with sulfidic lineages colored in yellow and nonsulfidic lineages in blue. Cacti 10 and 19 show clear clustering by ecotype. In cacti 1, 5, and 12, four of five sulfidic individuals cluster together.

### Convergent modifications of toxicity targets and detoxification pathways are evident at macroevolutionary scales

While selection on standing genetic variation and introgression can contribute to convergent evolution at microevolutionary scales, adaptive alleles are unlikely to be shared among lineages at macroevolutionary scales due to high phylogenetic and geographic distances separating gene pools (27). Absence of convergence in molecular mechanisms at broader phylogenetic scales might indicate the importance of contingency in evolution, as asserted by Gould (3). In contrast, the presence of convergence would indicate that fundamental constraints limit the number of solutions for a functional problem (28).

We used phylogenetic comparative analyses of gene expression and analyses of molecular evolution to detect patterns of molecular convergence in 10 lineages of sulfide spring poeciliids and ancestors in nonsulfidic habitats (Figure S1). This included members of five genera that span over 40 million years of divergence and occur in different biogeographic contexts (SI Appendix, Figure S1). We found evidence for convergence in both gene expression and sequence evolution. Variation in overall gene expression was strongly influenced by phylogenetic relationships (Figure 3A). However, 186 genes exhibited significant evidence for convergent expression shifts in sulfide spring fishes (Figure 3B; SI Appendix, Dataset S3), segregating lineages based on habitat type of origin, irrespective of phylogenetic relationships (Figure 3C). The only outlier was L*imia sulphurophila*, which clustered with nonsulfidic lineages despite significant expression differences with its sister, *L. perugiae*. Functional annotation indicated that genes with convergent expression shifts were enriched for biological processes associated with H_2_S detoxification (SQR pathway, Figure 3D), the processing of sulfur compounds, and H_2_S toxicity targets in OxPhos (SI Appendix, Figure S8, Table S14).

**Figure 3.**
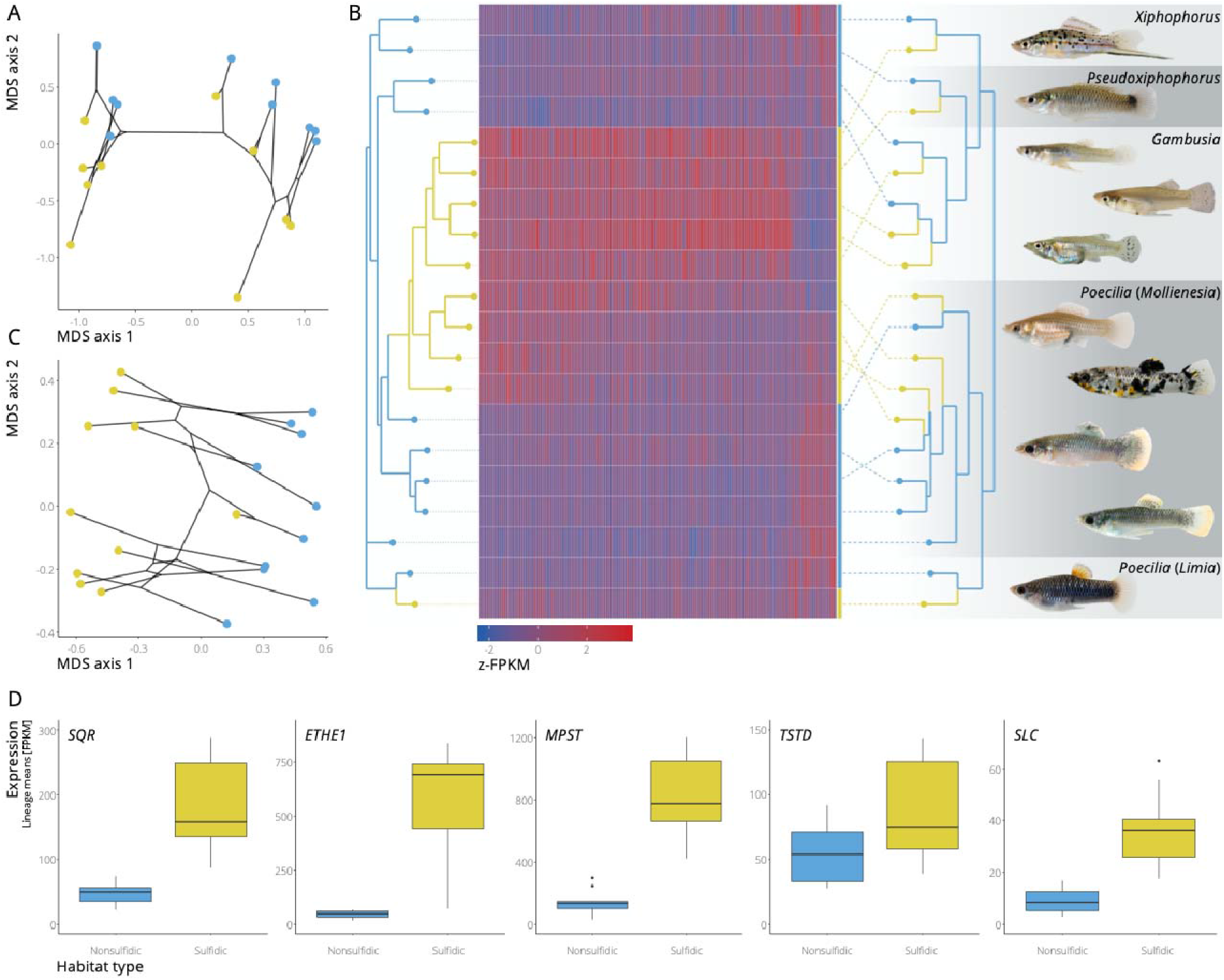
**A.** Multidimensional scaling (MDS) plot of overall gene expression patterns across 20 lineages of poeciliid fishes. Black lines represent phylogenetic relationships among lineages; color represents habitat type of origin (yellow: sulfidic; blue: nonsulfidic). **B.** Expression variation of 186 genes with evidence for convergent expression shifts (*z*-transformed FPKM, Fragments Per Kilobase of transcript per Million mapped reads). Colors represent expression levels as indicated by the scale. The neighbor-joining tree on the left groups species based on expression similarity. The cladogram on the right shows the phylogenetic relationship among lineages. Pictures on the side are examples of sulfide spring fishes (from top to bottom): *X. hellerii, P. bimaculatus, G. holbrooki, G. sexradiata, G. eurystoma, P. latipinna, P. sulphuraria* (Pich), *P. mexicana* (Tac), *P. mexicana* (Puy), and *L. sulphurophila*. **C.** MDS plot of the expression of 186 genes with evidence for convergent expression shifts. **D.** Boxplot with mean expression levels of different components of the SQR pathway across lineages from sulfidic (yellow) and nonsulfidic (blue) habitats.

We also identified 11 genes with elevated nonsynonymous to synonymous substitution rates across the phylogeny, including three mitochondrial genes that encode subunits of H_2_S’s toxicity target (*COX1* and *COX3*) and OxPhos complex III (*CYTB;* Dataset S4). Most amino acid substitutions in *COX1* and *COX3* occurred in a lineage-specific fashion, but convergent substitutions across clades occurred at six codons in *COX1* and two codons in *COX3* (Figure 4). These findings suggest that modifications of H_2_S toxicity targets and detoxification pathways are not only critical in the evolution of H_2_S tolerance in *P. mexicana*, but they have evolved in convergence in other lineages that were exposed to the same source of selection.

**Figure 4.**
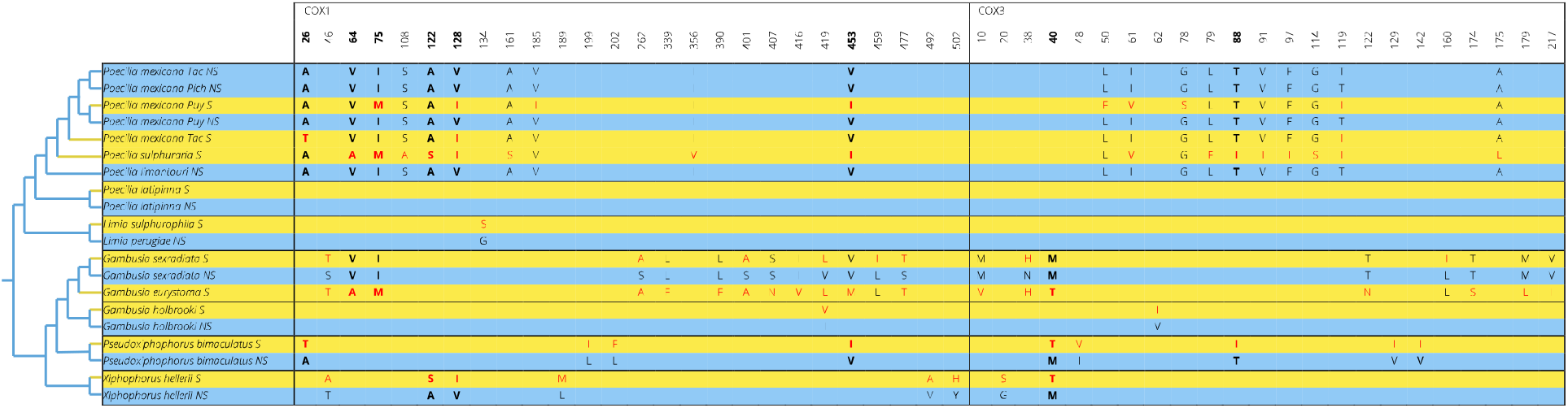
Amino acid differences in COX1 and COX3 between lineages from sulfidic (yellow) and nonsulfidic (blue) habitats. Derived amino acids are shown in red. Bold letters indicate codons with convergent amino acid substitutions in different clades (separated by black horizontal lines) of sulfide spring fishes.

## Conclusions

We capitalized on past evolutionary genetics studies that compared *P. mexicana* populations from sulfidic and nonsulfidic environments (10) to test hypotheses about functional ramifications of genetic differences and their impact on organismal performance. As predicted, we found that the repeated evolution of H_2_S tolerance in independent *P. mexicana* populations is mediated both by modifications of a direct toxicity target (causing increased resistance to H_2_S) and a pathway involved in detoxification (causing an increased ability to regulate mitochondrial H_2_S). Similar modifications to COX and SQR have been hypothesized to mediate H_2_S adaptation in other groups of organisms (29–31), but the evolutionary context and the consequences for mitochondrial function in these cases remained unknown. Overall, our analyses indicated that closely related populations can exhibit substantial differences in what we assume to be highly conserved physiological pathways associated with the function of mitochondria. Modification of mitochondrial processes consequently can be critical in mediating adaptation to different environmental conditions at microevolutionary scales, underscoring the long overlooked role of mitochondria in adaptive evolution (6).

Our comparative transcriptome analyses across a broader sampling of sulfide spring fishes further indicated that colonization of novel niches with extreme environmental conditions can arise through the convergent modification of conserved physiological pathways. The convergent evolution of high H_2_S tolerance across species is the result of repeated and predicted modifications of the same physiological pathways, genes, and—in some instances—codons associated with mitochondrial function. This convergence at multiple levels of biological organization is likely a consequence of constraint, because the explicit biochemical and physiological consequences of H_2_S limit the ways organisms can cope with its toxicity (32, 33). Due to these constraints, molecular convergence is not only evident at microevolutionary scales, where selection can repeatedly assemble related alleles into different genomic backgrounds, but also at macroevolutionary scales including lineages separated by over 40 million years of evolution.

That said, there is an inordinate amount of genetic and gene expression variation that seemingly varies idiosyncratically across different lineages. In comparative analyses of highly quantitative traits (such as H_2_S tolerance), there is an inherent bias to emphasize the importance of shared modifications in adaptation, while we tend to dismiss lineage-specific patterns as noise. But how lineage-specific genetic and gene expression variation interacts with molecular mechanisms that have evolved in convergence remains largely unknown for most study systems. So, if we replayed the tape of life, the same characters may make an appearance in the same setting, but the overall plots may still unfold in very different ways when many characters are part of the story.

## Methods

The following sections provide a synopsis of the procedures used in this study. Detailed materials and methods are provided in an SI Appendix.

### Sampling

Samples of *P. mexicana* for comparative biochemical and physiological analyses were collected from three population pairs from the Tacotalpa (Tac), Puyacatengo (Puy), and Pichucalco (Pich) drainages in Mexico, each including evolutionarily independent H_2_S-tolerant and ancestral, intolerant population (Table S1) (34). With the exception of measurements of mitochondrial H_2_S levels, which were conducted with common-garden-reared individuals, all assays were conducted with specimens collected in the field. For macroevolutionary analyses of convergence, we collected specimens from multiple species that have independently colonized H_2_S-rich springs in the United States, Mexico, and the island of Hispaniola, as well as from geographically and phylogenetically proximate lineages in nonsulfidic habitats. Our sampling included populations of *Poecilia latipinna* and *Gambusia holbrooki* in Florida; populations of the *Poecilia mexicana* species complex (including *Poecilia mexicana, Poecilia sulphuraria*, and *Poecilia limantouri*), the *Gambusia sexradiata* species complex (including *G. sexradiata* and *Gambusia eurystoma), Pseudoxiphophorus bimaculatus*, and *Xiphophorus hellerii* in Mexico; as well as populations of the L*imia perugiae* species complex (*L. perugiae* and L*imia sulphurophila*) in the Dominican Republic (see Table S1).

### Measurements of enzyme activity

Measurements of COX and SQR activities were conducted using isolated mitochondria from three *P. mexicana* population pairs and followed previously established methods. COX activity was quantified by measuring decrease in absorbance at 550 nm resulting from the protein’s oxidation of reduced cytochrome c (35, 36). Samples were assayed along a gradient of H_2_S concentrations between 0 and 0.9 μM. SQR activity was quantified by measuring the reduction of decyl-ubiquinone (37) along a gradient of H_2_S concentrations between 0 and 100 μM. Variation in enzyme activity for analyzed using linear mixed-effects models to estimate the effects of habitat type of origin (sulfidic vs. nonsulfidic), drainage, and H_2_S concentration. Alternative models were assessed by using Akaike Information Criteria with finite sample correction (AIC_C_) (38).

### Measurements of mitochondrial H_2_S

H_2_S exposure experiments and measurement of mitochondrial H_2_S were conducted using common-garden-raised F1 individuals from two population pairs (Tac and Puy). We used a mitochondria-specific H_2_S probe (MitoA) that can be injected into living organisms, where it accumulates within mitochondria due to its unique chemical structure (16). Inside mitochondria, MitoA reacts with H_2_S and is converted to MitoN. The ratio of MitoN/MitoA can be quantified using liquid chromatography with tandem mass spectrometry and serve as a metric of mitochondrial H_2_S. We used a MitoA protocol specifically validated for *P. mexicana* (39). Fish injected with MitoA were exposed to different environmental H_2_S concentrations for 5 hours. Gill, liver, brain, and muscle tissues were sampled from fish and use for the quantification of MitoN/MitoA ratios. Standardized MitoN/MitoA ratios were analyzed using linear mixed-effects models with habitat type of origin, drainage, and ambient H_2_S concentration as predictor variables. Alternative models were assessed by using Akaike Information Criteria with finite sample correction (AIC_C_).

### Measurements of mitochondrial function

Mitochondrial function was measured in mitochondria isolated from livers from three *P. mexicana* population pairs. Mitochondrial function was assayed using a Seahorse XFe96 Extracellular Flux Analyzer (Agilent Technologies, Santa Clara, CA, USA), which allows for the quantification of oxygen consumption rates (OCR) of isolated mitochondria in 96-well plates (40). Measuring mitochondrial OCR in presence of different substrates and inhibitors allows for the quantification of a variety of mitochondrial functions (41), and we measured basal respiration, maximal respiration, and spare respiratory capacity as indicators of mitochondrial function across a range of H_2_S concentrations between 0 and 90 μM. To compare responses in mitochondrial function to H_2_S between sulfidic and nonsulfidic populations, we employed a drug response analysis, where separate n-parameter logistic regressions were fit for each mitochondrial isolate with the metrics of mitochondrial function (basal respiration, maximal respiration, and spare capacity) as dependent variables and H_2_S concentration as independent variable (42). For each model, we quantified the area under the curve (AUC) based on Simpson’s rule, and higher AUC values represent an increased ability to maintain mitochondrial function in the presence of H_2_S. AUC values inferred for different mitochondrial isolates were then used as a dependent variable in linear models, with habitat type of origin and drainage as predictor variables. Alternative models were assessed by using Akaike Information Criteria with finite sample correction (AIC_C_).

### Comparative genomics and local ancestry analysis

To test hypotheses about the origin of putatively adaptive alleles, we re-analyzed data from Brown et al. (25), which included whole-genome sequences from all sites known to harbor H_2_S-tolerant populations of the *P. mexicana* species complex (*N*=5) as well as adjacent nonsulfidic habitats (*N*=5; Table S1). Raw reads were mapped to the *Xiphophorus maculatus* reference genome (43), and we called SNPs using Genome Analysis Toolkit (44). Local ancestry patters were identified with Saguaro (45), which uses a combination of a Hidden Markov Model and a self-organizing map to build ‘cacti’ (matrices of pairwise genetic distance between samples) that describe phylogenetic relationships among samples in specific genomic regions. Saguaro was executed for 29 iterations, which resulted in 30 cacti describing local ancestry patterns for segments of the genome.

### Comparative transcriptomics, phylogenetic comparative analyses, and molecular evolution

For comparative transcriptomic analyses, we isolated RNA from gills for 5-6 individuals each from 10 sulfidic and 10 non-sulfidic lineages distributed across multiple genera in the family Poeciliidae. Raw reads from transcriptome sequencing were mapped to the *Poecilia mexicana* reference genome (46), and the number of RNA-seq reads mapped to each gene was determined for each individual using cufflinks (47). To analyze patterns of gene expression across lineages, we used a phylogenetic comparative approach for analyzing gene expression variation to explicitly account for the effects of evolutionary relationships. Specifically, we used individual-level expression data of the top 5,000 expressed genes as dependent variables in Expression Variance and Evolution (EVE) models that identify genes exhibiting convergent shifts in expression upon the colonization of sulfidic habitats based on a phylogeny of focal taxa (58, 59). To identify genes with potential structural or functional changes in lineages from sulfidic habitats, we analyzed patterns of molecular evolution in protein coding genes included in the analysis of gene expression. For each gene, we generated a consensus sequences for all exons across individuals from the same populations and then used branch models implemented in the program codeml from the PAML package to test for evidence of positive selection (48).

## Acknowledgements

We thank the Centro de Investigación e Innovación para la Enseñanza y Aprendizaje (CIIEA), O. Cornejo, T. Morgan, D. Petrov, R. Rohlfs, and N. Rohner for their help. This work was supported by grants from the National Science Foundation (IOS-1463720, IOS-1557795, IOS-1557860, IOS-1931657 to JHS, JLK, and MT), the US Army Research Office (W911NF-15-1-0175, W911NF-16-1-0225 to JLK and MT), the Medical Research Council UK (MC_U105663142 to MPM), and by a Wellcome Trust Investigator Award (110159/Z/15/Z to MPM).

## Author contributions

Conceptualization: RG, NB, CH, JHS, JLK, MT; Funding: MPM, JHS, JLK, MT; Fieldwork: RG, NB, APB, LAR, CMRP, JLK, MT; Functional analyses: NB, CH, SA, GYL, MPM, LW, DL, JHS, MT; Genomics and transcriptomics: RG, APB, JLK, MT; Data analysis: RG, NB, CH, APB, JLK, MT; Writing original draft: RG, NB, JLK, MT; Writing, reviewing, editing: all authors.

## Data sharing

Data and code associated with biochemical and physiological analyses are available on GitHub (github.com/michitobler/convergent_h2s_evolution). All sequence data is available at NCBI under BioProject numbers PRJNA473350 and PRJNA608180.

