## Supplementary appendix for "Convergent Evolution of Conserved Mitochondrial Pathways Underlies Repeated Adaptation to Extreme Environments"

**This file includes:**

- Extended information on materials and methods
- Figures S1 to S9
- Tables S1 to S14

**Other Supplementary Materials for this manuscript include the following:**

- Dataset S1 as a separate Excel file
- Dataset S2 as a separate Excel file
- Dataset S3 as a separate Excel file
- Dataset S4 as a separate Excel file

### MATERIALS AND METHODS

#### 1. Background and hypotheses

The overall goals of our study were to test hypotheses about the mechanistic basis of $H_2S$ tolerance in evolutionarily replicated populations of *Poecilia mexicana* and about potential signatures of convergent evolution in sulfide spring fishes across the family Poeciliidae. *Poecilia mexicana* has been a model for the study of adaptation to $H_2S$-rich environments (1). This species is widely distributed in Mexico and Central America, inhabiting a wide range of freshwater habitats (2, 3). In southern Mexico, this species has also colonized $H_2S$-rich springs in four different river drainages of the Rio Grijalva basin (including the Rios Tacotalpa, Puyacatengo, Ixtapangajoya, and Pichucalco; see Figure S1). Colonization of springs has occurred independently, providing evolutionarily replicated population pairs in sulfidic and nonsulfidic habitats (4, 5). Sulfide spring populations are locally adapted and differ from ancestral populations in adjacent nonsulfidic habitats in a number of physiological, morphological, behavioral, and life-history traits (5-7). Most importantly, sulfide spring populations can tolerate $H_2S$ concentrations that are orders of magnitudes higher than what is considered lethal for most metazoans, while ancestral populations are susceptible to the toxic effects of $H_2S$ over short periods of time (5, 8), indicating strong survival selection that impacts fitness across small spatial scales. In addition, sulfide spring populations are reproductively isolated from adjacent populations in nonsulfidic water, with low levels of gene flow and high genetic differentiation despite small spatial scales (in some instances <100 m) and an absence of physical barriers that prevent fish movement (8). Reproductive isolation is in part mediated by natural and sexual selection against migrants (8, 9).

Comparative genomic and transcriptomic analyses of *P. mexicana* populations from sulfidic and nonsulfidic habitats have provided insights into mechanisms that potentially underlie tolerance to toxic levels of $H_2S$ (10-13). Collectively, these studies have indicated that modification of mitochondrial function may be critical in mediating adaptation to $H_2S$. Mitochondrial modifications may impact $H_2S$ tolerance through two mechanisms, resistance and regulation (14). Resistance involves modification of the direct toxicity target (cytochrome c oxidase, COX) of $H_2S$ and other components of the oxidative phosphorylation pathway (OxPhos). Regulation involves modification of physiological pathways associated with the maintenance of mitochondrial $H_2S$ concentrations; most notably the mitochondrial sulfide:quinone oxidoreductase (SQR) pathway. Past analyses have provided evidence for both resistance and regulation, as multiple components of the OxPhos and SQR pathways exhibit evidence for selection and/or heritable changes in gene regulation (10-13). However, the functional ramifications of amino acid substitutions and gene expression changes remain largely unknown.

Here, we hypothesized that the repeated modification of enzymes in the OxPhos and SQR pathways in *P. mexicana* populations from sulfidic habitats leads to an increased ability to maintain mitochondrial function in the presence of $H_2S$. We tested this hypothesis through multiple, complementary analyses:

1. Modification of COX—the primary toxicity target in OxPhos—is predicted to make the enzyme more resistant to the effects of $H_2S$. Previous analyses have already found evidence for this prediction in some *P. mexicana* populations (15). We followed up on this study and quantified COX activity across a broad range of $H_2S$ concentrations in multiple populations of *P. mexicana* from sulfidic and nonsulfidic habitats (see Section 2).
2. Modification of SQR—the first step in enzymatic $H_2S$ oxidation—is predicted to increase the rate of $H_2S$ detoxification. We tested this prediction by conducting enzyme activity assays, quantifying the rate at which $H_2S$ is oxidized by SQR in multiple populations of *P. mexicana* from sulfidic and nonsulfidic habitats (see Section 2).

3. If regulation of mitochondrial $H_2S$ is a primary mechanism of $H_2S$ tolerance, *P. mexicana* from sulfidic habitats should be able to maintain low endogenous $H_2S$ concentrations upon environmental exposure, whereas endogenous concentrations should increase in individuals from nonsulfidic habitats. We tested this prediction by experimentally exposing laboratory-reared individuals of *P. mexicana* to environmental $H_2S$ and quantifying endogenous $H_2S$ where it matters—in the mitochondria—using a novel mitochondria-targeted probe (MitoA) that can be deployed to measure $H_2S$ *in vivo* (see Section 3).
4. If regulation of endogenous $H_2S$ is a primary mechanism of $H_2S$ tolerance, mitochondrial function should be maintained in the presence of $H_2S$ even in sulfide spring populations that do not exhibit an $H_2S$-resistant COX. We tested this prediction by quantifying mitochondrial respiration across multiple *P. mexicana* populations in presence and absence of $H_2S$ *in vitro* (see Section 4).
5. We also asked whether adaptive alleles in *P. mexicana* from different sulfidic populations rose to high frequency through selection on standing genetic variation, or whether adaptive alleles arose independently in different sulfide spring populations through *de novo* mutations. To do so, we sequenced whole genomes of multiple *P. mexicana* populations to conduct local ancestry analyses (see Section 5).

Overall, our analyses in *P. mexicana* revealed that some sulfide spring populations have adaptive modifications to COX, an increased SQR activity, a higher capacity in maintaining low endogenous $H_2S$ concentration upon environmental exposure, and an ability to maintain mitochondrial respiration in presence of $H_2S$ irrespective of the presence of an $H_2S$-resistant toxicity target. Adaptive alleles at some loci (e.g., COX) arose independently in different populations, while there is also evidence for selection on standing genetic variation (e.g., genes associated with detoxification). Based on these findings, we asked whether other species that successfully colonized $H_2S$-rich habitats exhibit convergent modifications of genes associated with $H_2S$ toxicity and detoxification. This was possible because sulfide springs are widely distributed across the globe (16), and species of the family Poeciliidae are among the few metazoans that have colonized and adapted to these extreme environments (1). Collecting species from multiple genera allowed for the investigation of mechanisms underlying the evolution of $H_2S$ tolerance across broad phylogenetic scales (over 40 million years of divergence among lineages) and in different ecological and biogeographical contexts (Figure S1).

<u>We hypothesized that convergent molecular modifications underlie the convergent evolution of $H_2S$ tolerance across different lineages of poeciliid fishes</u>. We sequenced transcriptomes of five to six individuals from each of 10 independent lineages of sulfide spring fishes—as well as closely related lineages from nonsulfidic habitats—to test the following predictions (see Section 6):

6. Genes associated with $H_2S$ toxicity and detoxification should exhibit repeated changes of gene expression upon colonization of $H_2S$-rich habitats. We used phylogenetic comparative analyses to identify genes with convergent expression shifts in sulfide spring fishes.
7. Genes associated with $H_2S$ toxicity and detoxification should exhibit evidence for positive selection upon colonization of $H_2S$-rich habitats. We used analyses of molecular evolution to identify convergent targets of selection in sulfide spring fishes.

Note that all procedures used were approved by the Institutional Animal Care and Use Committees at Kansas State University (protocol #3418, 3473, 3581, 3886, 3992) and Oklahoma State University (protocol #1015, 1102). All statistical analyses were conducted in R version 3.6.1 (17) unless otherwise stated.

### 2. Enzyme activity assays

*2.1. Sample procurement*

We collected fish from three population pairs of *Poecilia mexicana* from the Tacotalpa (Tac), Puyacatengo (Puy), and Pichucalco (Pich) drainages in Mexico, each including an $H_2S$-tolerant and ancestral, intolerant population (Table S1). All fish were collected using a seine (2 × 5 m, 3 mm mesh size), immediately euthanized, and livers were extracted. Liver tissues were immediately frozen in liquid nitrogen and stored at -80° C upon return to Kansas State University.

*2.2. Cytochrome c oxidase (COX) activity*

2.2.1. Preparation of reagents and chemicals

All chemicals used to make the mitochondrial isolation buffer, incubation buffer, and substrates were purchased from Sigma-Aldrich (St. Louis, MO, USA), and solutions were made following the methods of previously published studies on OxPhos enzyme activity (18-20). The mitochondrial isolation buffer was prepared with 140 mM KCl, 10mM EDTA, 5mM $MgCl_2$, 20mM Hepes, and 2% bovine serum albumin (pH 7.3). Mitochondrial pellets were resuspended in an isotonic solution containing 100 mM KCl, 25 mM $K_2HPO_4$, and 5 mM $MgCl_2$ (pH 7.4). The reaction solution consisted of 25 mM $K_2HPO_4$, 5 mM $MgCl_2$, 2.5 mg/ml BSA, 0.6 mM lauryl maltoside, and 50 μM reduced cytochrome c (18). $H_2S$ was prepared with 1 mM $Na_2S \cdot 9\ H_2O$ and DDW purged of oxygen via bubbling of nitrogen under anoxic conditions. All solutions were prepared daily.

2.2.2. Isolation of mitochondria

For the isolation of mitochondria, 60 mg of liver tissue was placed directly into 500 μl of ice-cold mitochondrial isolation buffer and homogenized using a Dounce homogenizer (DWK Life Sciences, Millville, NJ, USA). An additional 500 μl of buffer was added to the sample prior to centrifugation. Homogenates were centrifuged at 600 g for 5 minutes at 4° C, and the supernatant was transferred to a new tube and centrifuged again at 9000 g for 5 minutes at 4° C. The pellet was resuspended in 500 μl of isotonic solution. 75 μl of the sample was taken to measure the total protein concentration (μg/ml) by using the BCA Assay reagent (Thermo Scientific Pierce, Rockford, IL, USA).

2.2.3. Measuring COX activity

COX activity was assayed using a 96-well plate spectrometer (BioTek Instruments Inc., Winooski, VT, USA), which allows for the measurement of change in absorbance of a reaction solution at different wavelengths and environmental parameters. Activity of COX is quantified following a decrease in absorbance at 550 nm resulting from the protein's oxidation of reduced cytochrome c (18, 19). Samples were assayed across a gradient of $H_2S$ concentrations between 0 and 0.9 μM in quadruplicate. Oxidation of cytochrome c was measured for 3 minutes at 11-second intervals and 25° C.

2.2.4. Statistical analyses

Variation in COX activity was analyzed using linear mixed-effects models as implemented in the LME4 package (21). Habitat type of origin (sulfidic or nonsulfidic) and drainage were used as factors, $H_2S$ concentration as a covariate, and sample ID was designated as a random effect. Alternative models were assessed by using Akaike Information Criteria with finite sample correction ($AIC_C$) (22). Besides the model with the lowest $AIC_C$ score, alternative models with $\Delta AIC_C < 2$ were considered equally supported (23).

*2.3. Sulfide:quinone oxireductase (SQR) activity*

2.3.1. Preparation of reagents and solutions

All chemicals used to make the mitochondrial isolation buffer, incubation buffer, and substrates were purchased from Sigma-Aldrich (St. Louis, MO, USA), and solutions were made following the methods of Hildebrandt and Grieshaber (20). The mitochondrial isolation buffer was prepared with 250 mM sucrose, 10 mM triethanolamine, 1 mM EGTA, 2 mM $K_2HPO_4$, 2 mM $KH_2PO_4$, and double distilled water (DDW) at pH 7.4. Incubation buffer was prepared with 250 mM sucrose, 10 mM triethanolamine, 1 mM EGTA, 2 mM $K_2HPO_4$, 2 mM $KH_2PO_4$, 5 mM $MgCl_2$, and DDW at pH 7.4. The reaction solution consisted of 20 mM Tris-HCl (pH 8.0) and the substrates 2 mM KCN, 100 μM decyl-ubiquinone, and 1 mM $H_2S$. $H_2S$ was prepared with 1 mM $Na_2S \cdot 9\ H_2O$ and DDW purged of oxygen via bubbling of nitrogen under anoxic conditions. Except for the $H_2S$ solution, which were prepared fresh daily, all solutions were stored at 4° C until used.

2.3.2. Isolation of mitochondria

To isolate mitochondria, 60 mg of liver tissue was placed directly into 500 μl ice-cold isolation buffer. Liver samples were homogenized on ice using a Dounce homogenizer (DWK Life Sciences, Millville, NJ, USA). Homogenates were centrifuged at 600 g for 10 minutes at 4° C, and the supernatant of each sample was again centrifuged at 9000 g for 20 minutes at 4° C. The resulting pellets were then resuspended in 500 ml ice-cold incubation buffer. 75 μl of the sample was taken to measure the total protein concentration (μg/ml) by using the BCA Assay reagent (Thermo Scientific Pierce, Rockford, IL, USA).

2.3.3. Measuring SQR activity

SQR activity was assayed using a 96-well plate spectrometer (BioTek Instruments Inc., Winooski, VT, USA). SQR activity was quantified following the reduction of decyl-ubiquinone. This reaction is cyanide-dependent in the absence of thioredoxin and sulfite, requiring the addition of KCN to the reaction mixture (24). Each reaction consisted of 2 mM KCN, 20 mM Tris-HCl, 100 μg/ml mitochondrial protein, decyl-ubiquinone, and $H_2S$. Samples were assayed across a gradient of $H_2S$ concentrations (0, 10, 30, 50, 80, and 100 μM) in duplicate. Reduction of decyl-ubiquinone was followed at 285 nm for 5 minutes using 50-second intervals at 25° C (20, 25).

2.3.4. Statistical analyses

Variation in SQR activity was analyzed using linear mixed-effects models as implemented in the LME4 package (21). Habitat type of origin (sulfidic or nonsulfidic) and drainage were used as factors, $H_2S$ concentration as a covariate, and sample ID was designated as a random effect. Alternative models were assessed by using Akaike Information Criteria with finite sample correction ($AIC_C$) (22). Besides the model with the lowest $AIC_C$ score, alternative models with $\Delta AIC_C < 2$ were considered equally supported (23).

### ***3. In vivo* measurement of endogenous $H_2S$ with MitoA**

*3.1. Sample procurement*

Fish used in this portion of the study were common-garden-raised F1 individuals from two population pairs in Mexico, Rios Puyacatengo (Puy) and Tacotalpa (Tac). Fish of the third population pair (Rio Pichucalco, Pich) were not available in the laboratory, because they are difficult to transport and maintain in aquaria. *Poecilia mexicana* were housed in 38-l tanks and maintained at 26° C with a 12h:12h light:dark cylce. Fish were fed *ad libitum* each day using commercial fish food and starved 24 h prior to experiments.

*3.2. Preparation of reagents and solutions*

The MitoA compounds (MitoA, MitoN, $d_{15}$-MitoA, and $d_{15}$-MitoN) used to quantify endogenous sulfide in this study were synthesized as previously described (26). A 10 mM stock solution of each MitoA compound was prepared with 200 proof ethanol. 60% acetonitrile was prepared with acetonitrile (Honeywell Research Chemicals, Mexico City, MX) and HPLC-grade water. $H_2S$ solutions (0, 1, 3, 4, and 5 mM) were prepared as described above.

*3.3. MitoA injection and $H_2S$ exposure*

To detect changes in endogenous $H_2S$ upon environmental exposure, we used MitoA, a mitochondrial specific probe that is capable of detecting $H_2S$ *in vivo* (26, 27). In brief, this probe is shown to readily accumulate within mitochondria due to its unique chemical structure, and upon binding to $H_2S$, it is converted to MitoN. MitoA and MitoN can then be quantified using tandem mass spectrometry, which allows for the analysis of change in $H_2S$ concentration within specific tissues and whole organisms (26). Methods described in the following section were specifically developed for *P. mexicana* (27).

On the day of each experiment, the 10 mM MitoA stock solution was diluted in phosphate-buffered saline (Sigma-Aldrich, St. Louis, MO, USA). Each fish was briefly anaesthetized using MS-222 (Sigma-Aldrich, St. Louis, MO, USA), restrained using a wet paper towel, and 8 nmol of MitoA in a volume of 50 µl was injected into the peritoneum using an insulin syringe with a 31-G needle (Becton, Dickinson and Company, Franklin Lakes, NJ, USA). Immediately after injection, each fish was placed into a 1-l container holding 250 mL of aerated water situated in a 25° C water bath and allowed to acclimate for 1 hour prior to the start of the trial. After acclimation, each container was sealed, and peristaltic pumps were used to provide a continuous supply of the designated $H_2S$ solution at a rate of 150 ml/h (28). Experimental exposures lasted 5 hours, when the final $H_2S$ concentrations were measured and fish were euthanized. Environmental $H_2S$ concentrations were measured using a methylene blue assay (Hach Company, Loveland, CO, USA). Gill, liver, brain, and muscle tissues were sampled from fish and flash frozen in liquid nitrogen. Tissue samples were stored at -80° C until further analysis.

*3.4. Extraction of MitoA and MitoN from tissues and LC-MS/MS analysis*

The methods for extraction of MitoA and MitoN followed previous protocols established for poeciliid fish (27). Tissues were weighed, placed into tubes containing 210 µl of a 60% acetonitrile solution spiked with internal standards, and underwent two separate homogenizations using a bead homogenizer. Between homogenizations, samples were centrifuged at 16000 g for 10 minutes, and supernatants were collected. The supernatant was incubated at 4° C for 30 minutes to precipitate proteins before being recentrifuged. Supernatants were then loaded onto 96-well Millipore Multiscreen filter plates (44 µm pore size) and centrifuged at 1100 g for 10 minutes. Filtrates were added to new collection tubes, dried using a SpeedVac vacuum concentrator, and stored until LC-MS/MS analysis.

LC-MS/MS analysis of tissue samples relied on the same methods used in Arndt et al. (26). Standard curves were generated using known amounts of MitoA and MitoN to calculate concentrations of each compound inside tissues, and the peak area of MitoA, MitoN, and standards were quantified using MASSLYNX version 4.1 (Waters Corporation, Milford, MA, USA).

*3.5. Statistical analyses*

Since we were interested in how endogenous $H_2S$ concentrations in different organs change upon environmental exposure, MitoN/MitoA ratios were standardized to the mean ratio of each organ and population. Variation in standardized MitoN/MitoA ratios was then analyzed using linear mixed-effects models as implemented in the LME4 package (21). Habitat type of origin (sulfidic or nonsulfidic), drainage, and $H_2S$ concentration were used as predictor variables. Organ type and individual ID were designated as random effects. Alternative models were assessed by using Akaike Information Criteria with finite sample correction ($AIC_C$) (22). Besides the model with the lowest $AIC_C$ score, alternative models with $\Delta AIC_C < 2$ were considered equally supported (23).

### 4. Mitochondrial Function

#### *4.1. Sample procurement*

This portion of the study focused on three sulfidic/nonsulfidic population pairs of *Poecilia mexicana* complex from the Rios Tacotalpa, Puyacatengo, and Pichucalco drainages in Mexico (Table S1). Fish were collected in their natural habitats using seines, transported to the laboratory at Oklahoma State University, and used for experiments within a few weeks of their capture.

#### *4.2. Preparation of reagents and solutions*

All chemicals that used to make mitochondrial isolation buffer (MSHE+BSA), mitochondrial assay solution (MAS, 3X) and substrates were purchased from Sigma-Aldrich (St. Louis, MO, USA). MSHE+BSA was prepared with 70 mM sucrose, 210 mM mannitol, 5 mM HEPES, 1 mM EGTA, 0.5% (w/v) fatty acid-free bovine serum albumin (BSA), and DDW. MAS, 3X was prepared with 210 mM sucrose, 660 mM mannitol, 30 mM $KH_2PO_4$, 15 mM $MgCl_2$, 6 mM HEPES, 3 mM EGTA, 0.6% (w/v) fatty acid free BSA, and DDW. 3X MAS was then diluted to make 1X MAS, which was used for the preparation of substrates, ADP, and respiration reagents. As substrates, 0.5 M succinate, 0.5 M malate, 0.5 M pyruvate, and 40 mM ADP were prepared with DDW. As respiration reagents, 10 μM oligomycin, 10 μM antimycin A/rotenone mix, and 3 μM FCCP [carbonyl cyanide 4-(trifluoromethoxy)-phenylhydrazone] (Seahorse XF Cell Mito Stress Test Kit, Santa Clara, CA, USA) were prepared with 1X MAS. All reagents and solutions were adjusted to pH 7.2 with potassium hydroxide. Except for the respiration reagents, which were prepared fresh on the day of each experiment, solutions were stored at 4° C until used.

#### *4.3. Isolation of mitochondria*

To isolate mitochondria, fish were euthanized and dissected to isolate livers, which were added to 500 μl MSHE+BSA and stored on ice. Due to the small size of the study species, livers from multiple individuals were pooled to obtain at least 60 mg of tissue, which provided a sufficient amount of mitochondria to run one coupling assay (see below). Liver samples were homogenized on ice with a Bio-Gen PRO200 (PRO Scientific, Oxford, CT, USA) for 10 seconds at the lowest speed, and then 500 μl MSHE+BSA were added to each homogenate. Homogenates were centrifuged at 600 g for 5 minutes at 4° C, and the filtered supernatant of each sample was again centrifuged at 5000 g for 5 minutes at 4° C. The resulting pellets were then resuspended in 1 ml 1X MAS. 100 μl of the sample was taken to measure the total protein concentration (mg/ml) by using the BCA Assay reagent (Thermo Scientific Pierce, Rockford, IL, USA).

#### *4.4. Mitochondrial coupling assay*

Mitochondrial function was assayed using a Seahorse XFe96 Extracellular Flux Analyzer (Agilent Technologies, Santa Clara, CA, USA), which allows for the quantification of oxygen consumption rates (OCR) of isolated mitochondria in 96-well plates (29). Energy demand and substrate availability for mitochondria can be tightly controlled in this setup through the sequential addition of compounds that either stimulate or inhibit components of the electron transport chain: the addition of ADP stimulates oxygen consumption, oligomycin blocks ATP synthase, FCCP uncouples oxygen consumption from ATP synthesis, and antimycin A and rotenone block mitochondrial complexes I and III, respectively. Measuring mitochondrial OCR in presence of these different compounds during a coupling assay allows for the quantification of a variety of mitochondrial functions (30), and we measured basal respiration, maximal respiration, and spare capacity as indicators of mitochondrial function across a range of $H_2S$ concentrations.

Prior to a coupling assay, the XFe96 sensor cartridge was pre-hydrated with calibrant solution (200 µl per well) overnight in a non-$CO_2$ 370c incubator. ADP, oligomycin, FCCP, and antimycin A/rotenone solutions (XF Cell Mito Stress Test Kit; Agilent Technologies) were then loaded into the four injection ports at the sensor cartridge for calibration. Meanwhile mitochondrial solution was added to each well of a 96-well plate (4 mg mitochondria per well) along with substrates (succinate, malate, and pyruvate) and 1X MAS. To test how mitochondrial function was affected by the presence of $H_2S$, sodium hydrosulfide (NaSH) was added directly to each well as $H_2S$ donor, just before a plate was placed in the analyzer. Overall, OCRs were measured at up to seven different doses of $H_2S$ (5, 15, 30, 50, 60, 80 and 90 µM) along with a nonsulfidic control (0 µM). After the addition of $H_2S$, coupling assays were conducted following the manufacturer's protocols. Each isolate was measured across a range of $H_2S$ concentrations, but tissue limitations prevented that all isolates were measured across all concentrations. All assays were run in triplicates, and OCR measurements were normalized by protein content.

*4.5. Statistical analyses*

To compare responses in mitochondrial function to $H_2S$ between sulfidic and nonsulfidic populations, we employed drug response analysis. Separate n-parameter logistic regressions were fit for each mitochondrial isolate with the metrics of mitochondrial function (basal respiration, maximal respiration, and spare capacity) as dependent variables and $H_2S$ concentration as independent variable, using the NPLR package (31). Based on regression models, AUC was estimated based on Simpson's rule, where higher AUC values represent an increased ability to maintain mitochondrial function in the presence of $H_2S$. AUC values inferred for different mitochondrial isolates were then used as a dependent variable in linear models using the STATS package (17). Alternative models were assessed by using Akaike Information Criteria with finite sample correction ($AIC_C$) (22). Besides the model with the lowest $AIC_C$ score, alternative models with $\Delta AIC_C$ <2 were considered equally supported (23).

**5. Comparative genomics and local ancestry analysis**

To test hypotheses about the origin of adaptive alleles, we re-analyzed data from Brown et al. (10) (NCBI short-read archive BioProject no. PRJNA473350) and used the raw reads from the guppy genome NCBI short-read archive accession no. SRR1171024 (32). Detailed methods about sample collection, library preparation, whole-genome sequencing, and single-nucleotide polymorphism (SNP) calling and filtering can be found there, and we provide a summary of general approaches here.

*5.1. Sample procurement*

We collected one individual each from all sites known to harbor $H_2S$-tolerant populations of the *P. mexicana* species complex (*N*=5) as well as adjacent nonsulfidic habitats (*N*=5; see Table S1 for details). Specimens were sacrificed on site, and muscle tissue was isolated and preserved in 96% ethanol.

*5.2. DNA extraction, library preparation, and sequencing*

A MagAttract High Molecular Weight DNA extraction kit was used to extract DNA, and the amount of extracted DNA was quantified with a Qubit fluorometer. Library preparation was completed using an Illumina's TruSeq DNA PCR-Free LT Library Preparation Kit. DNA was sheared using a Covaris M220 with a 550-bp insert size. Quantitative polymerase chain reaction (qPCR) with the KAPA Library Quantification kit for Illumina Sequencing Platforms was used for determining pooling amounts. Two equimolar pools of the libraries were created using the estimated concentrations from the qPCR. Libraries were then sequenced on an Illumina HiSeq 2500 platform at the Washington State University Genomics Core with paired-end, 100 bp runs.

*5.3. Mapping and variant calling*

Reads were mapped to the *Xiphophorus maculatus* reference genome [RefSeq accession number: GCF_000241075.1 (33)] using the BWA-MEM algorithm from BWA version 0.7.12-r1039 (34). SNPs were then called on a per-individual basis in the Genome Analysis Toolkit (GATK v. 3.5) with the EMIT_ALL_SITES option in the UnifiedGenotyper (35-37). Individual vcfs were merged with vcfmerge (vcftools version 0.1.15) (38). Filtering in GATK was conducted with the recommended filters QD < 2.0, FS > 60.0, MQ < 40.0, MQRankSum < -12.5, and ReadPosRankSum < -8.0. Additional filtering with vcftools retained only biallelic sites (--min-alleles 2 and --max-alleles 2), with at least 10 reads supporting the call (--minDP 10), quality scores greater than 30 (--minQ 30), and less than 10% missing data (--max-missing 0.9).

*5.4. Shared ancestry analysis*

To identify outliers between sulfidic and nonsulfidic populations, we first calculated $F_{ST}$ between the sulfidic and nonsulfidic samples, treating the two as separate groups. We also identified shared ancestry among lineages using Saguaro (39), which uses a combination of a Hidden Markov Model and a self-organizing map to build 'cacti' (matrices of pairwise genetic distance between samples) that describe phylogenetic relationships among samples in specific genomic regions. Saguaro builds an initial cactus (topology) based on the entire genome, then iteratively hypothesizes a new cactus for regions of the genome that do not fit the initial cactus. The process is iterated a user-defined number of times, and a new cactus is hypothesized in each iteration. To run Saguaro, our vcf was converted into the hidden Markov model format using the VCF2HMMFeature command within Saguaro (39). Saguaro was executed for 29 iterations, which resulted in 30 cacti describing local ancestry patterns for segments of the genome.

### 6. Comparative Transcriptomics

*6.1. Sample procurement*

We collected specimens from multiple lineages of poeciliid fishes that have independently colonized $H_2S$-rich springs in the United States, Mexico, and the island of Hispaniola, as well as from geographically and phylogenetically proximate lineages in nonsulfidic habitats. This approach led to sets of closely related lineages found in sulfidic and nonsulfidic habitats, including populations of *Poecilia latipinna* and *Gambusia holbrooki* in Florida; populations of the *Poecilia mexicana* species complex (including *Poecilia mexicana*, *Poecilia sulphuraria*, and *Poecilia limantouri*) (2), the *Gambusia sexradiata* species complex (including *G. sexradiata* and *Gambusia eurystoma*) (40), *Pseudoxiphophorus bimaculatus,* and *Xiphophorus hellerii* in Mexico; as well as populations of the *Limia perugiae* species complex (*L. perugiae* and *Limia sulphurophila*) (41) in the Dominican Republic (see Table S1 for localities and habitat associations of specific lineages). All fish were caught using a seine (2 × 5 meters; 3 mm mesh size). Immediately upon capture, adult females (*N*=5-6 per site; Table S1) were euthanized, and the gills were extracted from both sides of the body using sterilized scissors and forceps. Tissues were then preserved in RNAlater (Ambion Inc.) for subsequent analysis in the laboratory. We focused on gill tissues, because they are in direct contact with the toxic environment (14), mediate a variety of physiological processes involved in the maintenance of homeostasis (42, 43), and exhibit strong transcriptional responses upon exposure to $H_2S$ (13, 28). Note that data for some lineages of the *P. mexicana* species complex were reanalyzed from a previous study [GenBank BioProject accession number: PRJNA290391 (13)].

*6.2. RNA extraction, cDNA library preparation, and sequencing*

The general protocols for RNA extraction, library preparation and sequencing of all new samples followed procedures previously employed for *Poecilia* (13, 44). In brief, 10-30 mg of gill tissues from each individual were sealed in a Covaris TT1 TissueTube (Covaris, Inc., Woburn, MA, USA), frozen in liquid nitrogen, and pulverized. Total RNA was extracted from pulverized tissue using the NucleoSpin RNA kit (Machery-Nagel, Düren, Germany) following the manufacturer's protocol. mRNA isolation and cDNA library preparation were conducted using the NEBNext Poly(A) mRNA Magnetic Isolation Module (New England Biolabs, Inc., Ipswich, MA, USA) and NEBNext Ultra Directional RNA Library Prep Kit for Illumina (New England Biolabs, Inc., Ipswich, MA, USA) following the manufacturers' protocol with minor modifications. All cDNA libraries were individually barcoded, quantified using Qubit and an Agilent 2100 Bioanalyzer High Sensitivity DNA chip, and pooled in sets of 11-12 samples based on nanomolar concentrations. Samples were split into pools such that samples from different species and habitat types were sequenced together. There was no evidence for significant lane effects. Libraries were sequenced on an Illumina HiSeq 2500 with paired-end 101 base pair (bp) reads at the Washington State University Spokane Genomics Core. Due to low coverage, some samples were re-sequenced, and reads from separate runs were combined for each sample for data analysis.

*6.3. Mapping and annotation*

All raw RNA-seq reads were sorted by barcode and trimmed twice (quality 0 to remove adapters, followed by quality 24) using Trimgalore! version 0.4.0 (45). Trimmed reads for all 118 individuals were mapped to the *Poecilia mexicana* reference genome [RefSeq accession number: GCF_001443325.1 (46)] with an appended mitochondrial genome (GenBank Accession Number: KC992998.1) using BWA-MEM version 0.7.12 (47). An average of 92.6 ± 3.6% of reads mapped across all individuals (Table S13). We annotated genes from the *P. mexicana* reference genome by extracting the longest transcript per gene (using the perl script gff2fasta.pl[1]) and comparing them to the entries in the human SWISSPROT database (critical E-value 0.001; access date 04/15/2017) using BLASTx (48). We retained the top BLAST hit for each gene based on the top high-scoring segment pair.

*6.4. Phylogenetic analyses*

We used a phylogenetic framework established in a previous study (49) for subsequent analyses. In brief, we created lineage specific consensus sequences from 167 random genes, which were then aligned using ClustalW version 2.1 (50). Besides the focal lineages (Table S1), the alignment included sequences from *Fundulus heteroclitus* [GenBank accession number: JXMV00000000.1 (51)], which was used as an outgroup. We used jModelTest 2 version 2.1.10 (52) to determine the best partition scheme based on the most likely model of DNA substitution among 88 candidate models. Genes were then concatenated and partitioned according to the substitution models indicated by

[1] Available from https://github.com/ISUgenomics/common_scripts/blob/master/gff2fasta.pl

jModelTest and then used in Maximum Likelihood analyses using RAxML version 8.2.9 (53). We ran RAxML analyses under the Rapid Bootstrap Algorithm with 1,000 bootstrap replicates and used a thorough maximum likelihood (ML) search with each partition assigned its own GTR + $\Gamma$ + I model. These analyses recovered a robust hypothesis for evolutionary relationships among the focal taxa (bootstrap support generally ≥99%) that was consistent with previous phylogenetic studies of the family Poeciliidae (2, 54, 55). The only groups with moderately supported relationships included the two populations of *G. sexradiata* and *G. eurystoma* (bootstrap support 60%), which have previously been found to represent a complex of closely related and recently diverged species (40). The best scoring ML tree was utilized for subsequent analyses of gene expression variation and molecular evolution.

*6.5. Analysis of gene expression*

We used cufflinks version 2.2.1 (56) to quantify the number of RNA-seq reads mapped (measured as fragments per kilobase of transcript per million mapped reads, FPKM) to each gene in the *P. mexicana* reference genome for each individual. We then removed genes that did not have at least two FPKM in 70 or more individuals across all species. We subsequently focused our expression analyses on the remaining 5,000 genes with the highest average FPKM among all individuals. A total of 4,538 genes from this focal set (91%) had an annotation based on the human SWISSPROT database, including in 3,764 unique entries as some *P. mexicana* genes matched to the same human gene (see Datasets S3-4).

Since pairwise comparisons of gene expression across a broad phylogenetic sampling can be problematic (57), we used a phylogenetic comparative approach for analyzing gene expression variation to explicitly account for the effects of evolutionary relationships. Specifically, we used individual-level expression data as dependent variables in Expression Variance and Evolution (EVE) models that identify genes exhibiting convergent shifts in expression upon the colonization of sulfidic habitats based on the phylogenetic hypothesis described above (58, 59). EVE models implement an extended Ornstein-Uhlenbeck process that incorporates within-species expression variance to test for branch-specific shifts in gene expression by comparing likelihoods that an expression parameter ($\theta_i$) for a given gene is shared between two groups of lineages versus $\theta_i$ for that gene being significantly different between the groups (58). We designated branches associated with lineages from sulfidic habitats as one group and those associated with nonsulfidic lineages as another group and then contrasted $\theta_i$ for each gene between these two groups. For each gene, we employed a likelihood ratio test ($LRT_\theta$) contrasting the null hypothesis ($\theta_i^{sulfidic} = \theta_i^{nonsulfidic}$) to the alternative hypothesis ($\theta_i^{sulfidic} \neq \theta_i^{nonsulfidic}$) using a $X^2$ distribution to assess statistical significance (58). To account for multiple testing, we calculated FDR adjusted *P*-values using the Benjamini-Hochberg procedure (60). To explore the biological functions of genes with evidence for convergent expression shifts upon the colonization of sulfide springs (Dataset S3), we used a Gene Ontology (GO) enrichment analysis (61) as implemented in GOrilla [*p*-value threshold: 0.0001, accessed 03/07/2019 (62)]. The transcriptome wide annotation set served as the reference set, and a total of 3,711 unique SWISSPROT annotations were associated with a term in the GO database.

While we use gene expression variation to detect patterns of convergent evolution, we acknowledge that gene expression patterns are notoriously plastic, and organismal responses to physicochemical stressors occur at multiple hierarchical levels, including short-term acclimatization, developmental plasticity, genetic variation, and their interactions (63). Comparisons of gene expression between $H_2S$-tolerant and non-tolerant populations of *P. mexicana* have indicated a heritable basis underlying gene expression responses to $H_2S$ exposure, indicating that population differences are not a mere consequence of the environmental exposure history (12). However, our comparative data does not allow us to disentangle potential effects of plasticity in response to $H_2S$ exposure and genetic differences among sulfidic and nonsulfidic lineages. Even if mechanisms underlying gene expression variation differ among lineages, identifying sets of genes that are repeatedly recruited in $H_2S$ responses still provides insights into the role of convergence in $H_2S$ adaptation.

*6.6. Analyses of molecular evolution*

To identify genes with potential structural or functional changes in lineages from sulfidic habitats, we analyzed patterns of molecular evolution in 4,467 protein coding genes included in the analysis of gene expression. We first used the Genome Analysis Toolkit (GATK) version 3.5 (36) to realign mapped reads around indels (using the RealignerTargetCreator and IndelRealigner tools) and detect variants (using the UnifiedGenotyper tool) following GATK best practices (64, 65). Individuals were genotyped on a per-lineage basis with the EMIT_ALL_SITES output mode in the UnifiedGenotyper, creating a vcf file for each lineage. A custom pipeline was used to fill positions lacking sequence data in each vcf file with Ns. We used the setGT plugin of BCFtools version 1.3 (66) to designate the genotype at all variable sites as the major allele in each population vcf file. The coordinates for the coding sequences (CDS) were determined based on the *P. mexicana* gene annotation file (GTF) and subsequently used with the consensus tool in BCFtools to create population-specific consensus sequences across all individuals for each population. The consensus sequences for all exons of a gene were then concatenated, producing a single protein-coding sequence for each gene and lineage. The nucleotide sequences for all populations were aligned using the corresponding amino acid sequence with translatorX (67), which implements the MAFFT progressive alignment method (68), ultimately resulting in an in-frame multispecies alignment for each gene. Phylogenetic trees were constructed for each gene using a maximum likelihood approach implemented with the DNAML program in PHYLIP version 3.696 (69).

To identify genes under positive selection in sulfide spring lineages, we used branch models implemented in the program codeml from the PAML package version 4.9a (70). Branch models in codeml estimate variation in the rates of synonymous to synonymous substitutions (ω) for each codon in an alignment across branches in a phylogeny. We supplied codeml with the in-frame multispecies alignment and maximum likelihood tree (gene tree) for each of our focal genes, designating lineages from sulfidic habitats as foreground branches and lineages from nonsulfidic habitats as background branches in the tree file to test for shifts in ω between foreground and background branches. Specifically, we compared a two-ratio branch model, calculating separate ω values for the foreground and background groups (*i.e.*, sulfidic and nonsulfidic lineages) designated in the tree file (M2; model = 2, NSsites = 0), against a null model with a single ω estimated for the tree (M0; model = 0, NSsites = 0) for each gene. We assessed significance using a likelihood-ratio test, calculating the test statistic as two times the difference between the log-likelihood of the branch model and the-log likelihood of the null model, with *P*-values calculated from a $X^2$ approximation (71). To account for multiple testing, we calculated FDR adjusted *P*-values using the Benjamini-Hochberg procedure (see above). Genes with evidence for positive selection across sulfidic lineages were used in a GO enrichment analysis, as described above (see Gene expression analysis).

**Table S1.** List of lineages from sulfidic and nonsulfidic habitats included in this study. The table provides descriptions of the collection localities, information about the presence or absence of $H_2S$, latitude and longitude (Lat/Long), as well as the sample sizes for the quantification of SQR activity ($N_{SQR}$; number of biological replicates, each tested across multiple concentrations), endogenous $H_2S$ concentrations ($N_{mitoA}$; number of individuals with multiple tissues analyzed for each), mitochondrial function ($N_{resp}$; number of biological replicates, each tested across multiple concentrations), comparative genomics ($N_{genomics}$), as well as transcriptomics ($N_{RNAseq}$; number of individuals).

| ID | Species | Locality | $H_2S$ | Lat/Long | $N_{SQR}$ | $N_{mitoA}$ | $N_{resp}$ | $N_{genomics}$ | $N_{RNAseq}$ |
|---|---|---|---|---|---|---|---|---|---|
| 1 | *Poecilia mexicana* | Arroyo Bonita, Rio Tacotalpa drainage, Tabasco, MX | – | 17.427/-92.752 | 18 | 14 | 5 | 1 | 6 |
| 2 | *Poecilia mexicana* | Arroyo Rosita, Rio Pichucalco drainage, Chiapas, MX | – | 17.485/-93.104 | 12 | | 3 | 1 | 6 |
| 3 | *Poecilia mexicana* | La Lluvia springs, Rio Puyacatengo drainage, Tabasco, MX | + | 17.464/-92.895 | 18 | 16 | 4 | 1 | 5 |
| 4 | *Poecilia mexicana* | Rio Puyacatengo, Rio Puyacatengo drainage, Tabasco, MX | – | 17.510/-92.914 | 12 | 16 | 4 | 1 | 6 |
| 5 | *Poecilia mexicana* | El Azufre, Rio Tacotalpa drainage, Tabasco, MX | + | 17.438/-92.775 | 24 | 14 | 4 | 1 | 5 |
| 6 | *Poecilia sulphuraria* | Baños del Azufre, Rio Pichucalco drainage, Tabasco, MX | + | 17.552/-92.999 | 24 | | 2 | 1 | 6 |
| 7 | *Poecilia mexicana* | Rio El Azufre, west branch, Rio Pichucalco drainage, Chiapas, MX | - | 17.556/-93.008 | | | | 1 | |
| 8 | *Poecilia mexicana* | Rio Ixtapangajoya, Rio Ixtapangajoya drainage, Chiapas, MX | - | 17.495/-92.998 | | | | 1 | |
| 9 | *Poecilia sulphuraria* | La Gloria springs, Rio Pichucalco drainage, Chiapas, MX | + | 17.532/-93.015 | | | | 1 | |
| 10 | *Poecilia thermalis* | La Esperanza springs, Rio Ixtapangajoya drainage, Tabasco, MX | + | 17.511/-92.983 | | | | 1 | |
| 11 | *Poecilia limantouri* | Rio Garces, Rio Panuco drainage, Hidalgo, MX | – | 20.940/-98.282 | | | | | 6 |
| 12 | *Poecilia latipinna* | Green Springs, St. Johns River drainage, Florida, USA | + | 28.863/-81.248 | | | | | 6 |
| 13 | *Poecilia latipinna* | Mariner's Cove, Lake Monroe, St. Johns River drainage, Florida, USA | – | 28.857/-81.239 | | | | | 6 |
| 14 | *Limia sulphurophila* | Balnearios La Zurza, Lago Enriquillo basin, Independencia, DR | + | 18.398/-71.570 | | | | | 6 |
| 15 | *Limia perugiae* | Stream in Cabral, Rio Yaque del Sur drainage, Barahona, DR | – | 18.246/-71.223 | | | | | 6 |
| 16 | *Gambusia sexradiata* | Mogote del Puyacatengo, Rio Puyacatengo drainage, Tabasco, MX | + | 17.582/-92.900 | | | | | 6 |
| 17 | *Gambusia sexradiata* | Laguna Sitio Grande, Rio Ixtapangajoya drainage, Tabasco, MX | – | 17.677/-92.997 | | | | | 6 |
| 18 | *Gambusia eurystoma* | Baños del Azufre, Rio Pichucalco drainage, Tabasco, MX | + | 17.552/-92.999 | | | | | 6 |
| 19 | *Gambusia holbrooki* | Green Springs, St. Johns River drainage, Florida, USA | + | 28.863/-81.248 | | | | | 6 |
| 20 | *Gambusia holbrooki* | Mariner's Cove, Lake Monroe, St. Johns River drainage, Florida, USA | – | 28.857/-81.239 | | | | | 6 |

| | | | | | |
|---|---|---|---|---|---|
| 21 | *Pseudoxiphophorus bimaculatus* | La Gloria springs, Rio Pichucalco drainage, Chiapas, MX | + | 17.532/-93.015 | 6 |
| 22 | *Pseudoxiphophorus bimaculatus* | Arroyo Pujil, Rio Ixtapangajoya drainage, Chiapas, MX | – | 17.476/-92.986 | 6 |
| 23 | *Xiphophorus hellerii* | La Gloria springs, Rio Pichucalco drainage, Chiapas, MX | + | 17.532/-93.015 | 6 |
| 24 | *Xiphophorus hellerii* | Rio El Azufre, west branch, Rio Pichucalco drainage, Chiapas, MX | – | 17.556/-93.008 | 6 |

**Table S2.** Results of mixed-effect linear models analyzing variation in COX activity. Models are ordered based on $\Delta AIC_C$ values. The null model included biological replicate ID as a random factor. Only one model (bold) exhibited $\Delta AIC_C < 2$.

| Model | Terms | AICc | ΔAICc | df | Weight |
|---|---|---|---|---|---|
| **m8** | **Habitat × $H_2S$ + (1\|ID)** | **-30.8** | **0.0** | **6** | **0.988** |
| m5 | Habitat + $H_2S$ + (1\|ID) | -21.0 | 9.8 | 5 | 0.007 |
| m11 | Habitat × Drainage × $H_2S$ + (1\|ID) | -18.9 | 11.9 | 14 | 0.003 |
| m3 | $H_2S$ + (1\|ID) | -18.6 | 12.3 | 4 | 0.002 |
| m10 | Habitat + Drainage + $H_2S$ + (1\|ID) | -11.8 | 19.0 | 7 | <0.001 |
| m6 | Drainage + $H_2S$ + (1\|ID) | -10.0 | 20.9 | 6 | <0.001 |
| m9 | Drainage × $H_2S$ + (1\|ID) | -5.3 | 25.5 | 8 | <0.001 |
| m1 | Habitat + (1\|ID) | 45.1 | 76.0 | 4 | <0.001 |
| null | (1\|ID) | 47.6 | 78.4 | 3 | <0.001 |
| m4 | Habitat + Drainage + (1\|ID) | 54.2 | 85.1 | 6 | <0.001 |
| m2 | Drainage + (1\|ID) | 56.1 | 87.0 | 5 | <0.001 |
| m7 | Habitat × Drainage + (1\|ID) | 58.6 | 89.4 | 8 | <0.001 |

**Table S3.** Results of the top mixed-effect linear model analyzing variation in COX activity (m8, Table S2).

| Predictors | Estimates | CI | *P* |
|---|---|---|---|
| (Intercept) | 0.8698 | 0.7567 – 0.9830 | <0.001 |
| Habitat | 0.0749 | -0.0887 – 0.2385 | 0.369 |
| $H_2S$ | -0.6545 | -0.7774 – -0.5317 | <0.001 |
| Habitat × $H_2S$ | 0.3612 | 0.1836 – 0.5388 | <0.001 |
| Random Effects | | | |
| $\sigma^2$ | 0.03 | | |
| $\tau_{00\text{-}ID}$ | 0.03 | | |
| ICC | 0.49 | | |
| $N_{ID}$ | 23 | | |
| Observations | 138 | | |
| Marginal $R^2$ | 0.417 | | |
| Conditional $R^2$ | 0.700 | | |

**Table S4.** Results of mixed-effect linear models analyzing variation in SQR activity. Models are ordered based on $\Delta AIC_C$ values. The null model included biological replicate ID as a random factor. Only one model (bold) exhibited $\Delta AIC_C < 2$.

| Model | Terms | AICc | ΔAICc | df | Weight |
|---|---|---|---|---|---|
| **m8** | **Habitat × $H_2S$ + (1\|ID)** | **-817.8** | **0.0** | **6** | **0.752** |
| m1 | Habitat + (1\|ID) | -815.5 | 2.3 | 4 | 0.239 |
| null | (1\|ID) | -808.8 | 9.0 | 3 | 0.009 |
| m5 | Habitat + $H_2S$ + (1\|ID) | -795.5 | 22.3 | 5 | <0.001 |
| m4 | Habitat + Drainage + (1\|ID) | -789.0 | 28.8 | 6 | <0.001 |
| m3 | $H_2S$ + (1\|ID) | -788.7 | 29.1 | 4 | <0.001 |
| m2 | Drainage + (1\|ID) | -784.4 | 33.3 | 5 | <0.001 |
| m10 | Habitat + Drainage + $H_2S$ + (1\|ID) | -768.9 | 48.9 | 7 | <0.001 |
| m7 | Habitat × Drainage + (1\|ID) | -766.7 | 51.1 | 8 | <0.001 |
| m6 | Drainage + $H_2S$ + (1\|ID) | -764.2 | 53.6 | 6 | <0.001 |
| m9 | Drainage × $H_2S$ + (1\|ID) | -727.3 | 90.4 | 8 | <0.001 |
| m11 | Habitat × Drainage × $H_2S$ + (1\|ID) | -706.4 | 111.4 | 14 | <0.001 |

**Table S5.** Results of the top mixed-effect linear model analyzing variation in SQR activity (m8, Table S4).

| Predictors | Estimates | CI | *P* |
|---|---|---|---|
| (Intercept) | 0.0019 | -0.0001 – 0.0040 | 0.060 |
| Habitat | -0.0011 | -0.0037 – 0.0015 | 0.421 |
| $H_2S$ | -0.0001 | -0.0001 – 0.0000 | <0.001 |
| Habitat × $H_2S$ | 0.0002 | 0.0001 – 0.0002 | <0.001 |
| **Random Effects** | | | |
| $\sigma^2$ | 0.00 | | |
| $\tau_{00\text{-}ID}$ | 0.00 | | |
| ICC | 0.10 | | |
| $N_{ID}$ | 19 | | |
| Observations | 108 | | |
| Marginal $R^2$ | 0.513 | | |
| Conditional $R^2$ | 0.563 | | |

**Table S6.** Results of mixed-effect linear models analyzing variation in relative mitochondrial $H_2S$ concentrations (MitoN/MitoA). Models are ordered based on $\Delta AIC_C$ values. The null model included individual ID and organ type as random factors. Only one model (bold) exhibited $\Delta AIC_C$ < 2.

| Model | Terms | AICc | ΔAICc | df | Weight |
|---|---|---|---|---|---|
| **m4** | **$H_2S$ + Habitat + (1\|ID) + (1\|Organ)** | **510.1** | **0.0** | **6** | **0.594** |
| m2 | Habitat + (1\|ID) + (1\|Organ) | 513.7 | 3.6 | 5 | 0.100 |
| m7 | $H_2S$ × Habitat + (1\|ID) + (1\|Organ) | 514.0 | 3.9 | 7 | 0.086 |
| m9 | Habitat × Drainage + (1\|ID) + (1\|Organ) | 514.2 | 4.1 | 7 | 0.076 |
| m10 | $H_2S$ + Habitat + Drainage + (1\|ID) + (1\|Organ) | 514.4 | 4.3 | 7 | 0.069 |
| m11 | $H_2S$ × Habitat × Drainage + (1\|ID) + (1\|Organ) | 514.8 | 4.7 | 11 | 0.058 |
| m6 | Habitat + Drainage + (1\|ID) + (1\|Organ) | 517.9 | 7.8 | 6 | 0.012 |
| m1 | $H_2S$ + (1\|ID) + (1\|Organ) | 519.8 | 9.7 | 5 | 0.005 |
| null | (1\|ID) + (1\| Organ) | 522.9 | 12.8 | 4 | 0.001 |
| m5 | $H_2S$ + Drainage + (1\|ID) + (1\|Organ) | 523.8 | 13.7 | 6 | <0.001 |
| m3 | Drainage + (1\|ID) + (1\|Organ) | 526.8 | 16.7 | 5 | <0.001 |
| m8 | $H_2S$ × Drainage + (1\|ID) + (1\|Organ) | 528.0 | 17.9 | 7 | <0.001 |

**Table S7.** Results of the top mixed-effect linear model analyzing variation in relative endogenous $H_2S$ concentrations activity (m4, Table S6).

| Predictors | Estimates | CI | *P* |
|---|---|---|---|
| (Intercept) | 1.2580 | 0.9922 – 1.5238 | **<0.001** |
| $H_2S$ | 0.1812 | 0.0706 – 0.2917 | **0.001** |
| Habitat | -0.5064 | -0.7563 – -0.2566 | **<0.001** |
| **Random Effects** | | | |
| $\sigma^2$ | 0.48 | | |
| $\tau_{00\text{-}ID}$ | 0.11 | | |
| $\tau_{00\text{-}Organ}$ | 0.01 | | |
| ICC | 0.19 | | |
| $N_{ID}$ | 60 | | |
| $N_{Organ}$ | 4 | | |
| Observations | 216 | | |
| Marginal $R^2$ | 0.160 | | |
| Conditional $R^2$ | 0.318 | | |

**Table S8.** Results of linear models analyzing variation in basal mitochondrial respiration. Models are ordered based on $\Delta AIC_C$ values. The null model only included the intercept. Even through m2 exhibited $\Delta AIC_C < 2$, the null model was best supported overall.

| Model | Terms | AICc | ΔAICc | df | Weight |
|---|---|---|---|---|---|
| **null** | **1** | **180.2** | **0.0** | **2** | **0.453** |
| **m2** | **1 + Habitat** | **181.4** | **1.2** | **3** | **0.248** |
| m1 | 1 + Drainage | 182.4 | 2.3 | 4 | 0.146 |
| m3 | 1 + Drainage + Habitat | 183.4 | 3.2 | 5 | 0.091 |
| m4 | 1 + Drainage × Habitat | 184.2 | 4.0 | 7 | 0.062 |

**Table S9.** Results of linear models analyzing variation in maximal mitochondrial respiration. Models are ordered based on $\Delta AIC_C$ values. The null model only included the intercept. Only one model (bold) exhibited $\Delta AIC_C < 2$.

| Model | Terms | AICc | ΔAICc | df | Weight |
|---|---|---|---|---|---|
| **m4** | **1 + Drainage × Habitat** | **148.8** | **0.0** | **7** | **0.996** |
| m3 | 1 + Drainage + Habitat | 159.9 | 11.1 | 5 | 0.004 |
| m1 | 1 + Drainage | 188.7 | 39.9 | 4 | <0.001 |
| m2 | 1 + Habitat | 191.1 | 42.3 | 3 | <0.001 |
| null | 1 | 198.1 | 49.3 | 2 | <0.001 |

**Table S10.** Results of the top linear model analyzing variation in maximal mitochondrial respiration (m4, Table S9).

| Factor | SS | df | *F* | *P* |
|---|---|---|---|---|
| (Intercept) | 333.1 | 1 | 13.015 | 0.002 |
| Drainage | 2803.7 | 2 | 54.782 | <0.001 |
| Habitat | 2313.6 | 1 | 90.412 | <0.001 |
| Drainage × Habitat | 578.2 | 2 | 11.298 | 0.001 |
| Residuals | 409.4 | 16 | | |

**Table S11.** Results of linear models analyzing variation in spare respiratory capacity. Models are ordered based on $\Delta AIC_C$ values. The null model only included the intercept. Only one model (bold) exhibited $\Delta AIC_C < 2$.

| Model | Terms | AICc | ΔAICc | df | Weight |
|---|---|---|---|---|---|
| **m4** | **1 + Drainage × Habitat** | **167.5** | **0.0** | **7** | **0.870** |
| m3 | 1 + Drainage + Habitat | 171.3 | 3.8 | 5 | 0.130 |
| m1 | 1 + Drainage | 189.3 | 21.8 | 4 | <0.001 |
| m2 | 1 + Habitat | 190.9 | 23.4 | 3 | <0.001 |
| null | 1 | 196.6 | 29.0 | 2 | <0.001 |

**Table S12.** Results of the top linear model analyzing variation in spare respiratory capacity (m4, Table S11).

| Factor | SS | df | *F* | *P* |
|---|---|---|---|---|
| (Intercept) | 273.7 | 1 | 4.552 | 0.049 |
| Drainage | 2808.1 | 2 | 23.351 | <0.001 |
| Habitat | 2249.4 | 1 | 37.410 | <0.001 |
| Drainage × Habitat | 698.1 | 2 | 5.805 | 0.013 |
| Residuals | 962.0 | 16 | | |

**Table S13.** Descriptive statistics for Illumina sequencing reads used for comparative transcriptome analyses of poeciliids from sulfidic and nonsulfidic environments. For each lineage, the table lists the average number of reads per individual after trimming for quality, the average number of reads that mapped to the reference genome, and the average proportion of reads that mapped to the reference genome. Error margins are provided in the form of standard errors. Lineages included are numbered corresponding to Table S1.

| ID | Species | $H_2S$ | N | Reads | Mapped reads | Percent mapped |
|---|---|---|---|---|---|---|
| 1 | *Poecilia mexicana* Tac | – | 6 | 13,179,072 ± 782,269 | 12,615,832 ± 758,991 | 0.957 ± 0.004 |
| 2 | *Poecilia mexicana* Pich | – | 6 | 11,945,216 ± 1,160,788 | 11,296,904 ± 1,105,644 | 0.945 ± 0.005 |
| 3 | *Poecilia mexicana* Puy | + | 5 | 20,161,962 ± 2,212,689 | 18,735,621 ± 2,096,062 | 0.928 ± 0.007 |
| 4 | *Poecilia mexicana* Puy | – | 6 | 19,700,214 ± 1,865,276 | 18,355,384 ± 1,774,642 | 0.931 ± 0.007 |
| 5 | *Poecilia mexicana* Tac | + | 6 | 14,400,204 ± 2,690,186 | 13,807,309 ± 2,592,441 | 0.958 ± 0.003 |
| 6 | *Poecilia sulphuraria* Pich | + | 6 | 12,267,262 ± 1,644,534 | 11,740,414 ± 1,577,010 | 0.957 ± 0.002 |
| 11 | *Poecilia limantouri* | – | 6 | 26,586,625 ± 9,141,136 | 25,188,316 ± 8,720,553 | 0.946 ± 0.004 |
| 12 | *Poecilia latipinna* | + | 6 | 43,960,656 ± 7,696,566 | 42,661,324 ± 7,443,424 | 0.971 ± 0.001 |
| 13 | *Poecilia latipinna* | – | 6 | 28,591,830 ± 2,092,989 | 27,606,064 ± 2,013,339 | 0.966 ± 0.001 |
| 14 | *Limia sulphurophila* | + | 6 | 26,523,621 ± 3,993,393 | 25,311,603 ± 3,807,545 | 0.955 ± 0.002 |
| 15 | *Limia perugiae* | – | 6 | 26,579,863 ± 3,206,613 | 25,109,831 ± 3,062,313 | 0.944 ± 0.003 |
| 16 | *Gambusia sexradiata* | + | 6 | 31,896,270 ± 4,598,187 | 28,595,855 ± 4,024,693 | 0.900 ± 0.010 |
| 17 | *Gambusia sexradiata* | – | 6 | 35,598,499 ± 0,503,441 | 29,867,961 ± 8,007,416 | 0.874 ± 0.028 |
| 18 | *Gambusia eurystoma* | + | 6 | 19,514,414 ± 4,021,075 | 16,883,905 ± 3,364,767 | 0.870 ± 0.006 |
| 19 | *Gambusia holbrooki* | + | 6 | 29,612,503 ± 5,887,338 | 26,944,912 ± 5,401,728 | 0.909 ± 0.003 |
| 20 | *Gambusia holbrooki* | – | 6 | 35,914,764 ± 8,032,747 | 32,308,042 ± 7,135,952 | 0.903 ± 0.004 |
| 21 | *Pseudoxiphophorus bimaculatus* | + | 6 | 39,545,078 ± 6,735,006 | 36,031,145 ± 6,142,661 | 0.910 ± 0.002 |
| 22 | *Pseudoxiphophorus bimaculatus* | – | 6 | 35,243,934 ± 7,309,241 | 32,539,388 ± 6,787,986 | 0.922 ± 0.001 |
| 23 | *Xiphophorus hellerii* | + | 6 | 29,296,035 ± 5,042,914 | 26,660,330 ± 4,574,585 | 0.911 ± 0.004 |
| 24 | *Xiphophorus hellerii* | – | 6 | 30,761,274 ± 6,214,700 | 28,144,578 ± 5,721,047 | 0.914 ± 0.003 |

**Table S14.** Gene Ontology terms for genes exhibiting convergent shifts toward higher expression in sulfide spring fishes that had evidence for significant enrichment (after FDR correction). Provided are the GO term identification number, a description of each GO term, *P*-value associated with the enrichment, false-discovery rate (*q*), the level of enrichment, the total number of genes in the reference set (*N*), the total number of genes with a specific GO term in the reference set (*B*), the number of genes in the target set (*n*), and the number of genes in the intersection (*b*). We also provide a list of all upregulated genes associated with each GO term. Note that there was no evidence for enrichment in genes exhibiting convergent shifts toward lower expression in sulfide spring fishes. Enrichment of upregulated genes included biological processes associated with sulfide detoxification as well as sulfur metabolism and transport, mitochondrial function and energy metabolism, as well as protein synthesis and disassembly, which are indirectly affected by the presence of $H_2S$ (14).

| GO Term | Description | *P* | *q* | Enrichment | *N* | *B* | *n* | *b* | Genes |
|---|---|---|---|---|---|---|---|---|---|
| GO:0070221 | Sulfide oxidation; using sulfide:quinone oxidoreductase | 2.30E-06 | 2.00E-03 | 25.43 | 3713 | 4 | 146 | 4 | SQRDL: sulfide quinone reductase-like (yeast); SLC25A10: solute carrier family 25 (mt carrier; dicarboxylate transporter); member 10; ETHE1: ethylmalonic encephalopathy 1; TSTD1: thiosulfate sulfurtransferase (rhodanese)-like domain containing 1 |
| GO:0019418 | Sulfide oxidation | 2.30E-06 | 2.18E-03 | 25.43 | 3713 | 4 | 146 | 4 | SQRDL: sulfide quinone reductase-like (yeast); SLC25A10: solute carrier family 25 (mt carrier; dicarboxylate transporter); member 10; TSTD1: thiosulfate sulfurtransferase (rhodanese)-like domain containing 1; ETHE1: ethylmalonic encephalopathy 1 |
| GO:0070813 | Hydrogen sulfide metabolic process | 1.11E-05 | 7.76E-03 | 20.35 | 3713 | 5 | 146 | 4 | SQRDL: sulfide quinone reductase-like (yeast); CBS: cystathionine-beta-synthase; MPST: mercaptopyruvate sulfurtransferase; ETHE1: ethylmalonic encephalopathy 1 |
| GO:0000098 | Sulfur amino acid catabolic process | 7.32E-05 | 3.83E-02 | 14.53 | 3713 | 7 | 146 | 4 | CDO1: cysteine dioxygenase type 1; GADL1: glutamate decarboxylase-like 1; CBS: cystathionine-beta-synthase; MPST: mercaptopyruvate sulfurtransferase |
| GO:0006534 | Cysteine metabolic process | 9.74E-06 | 7.27E-03 | 14.13 | 3713 | 9 | 146 | 5 | CDO1: cysteine dioxygenase type 1; GCLM: glutamate-cysteine ligase; modifier subunit; CBS: cystathionine-beta-synthase; GCLC: glutamate-cysteine ligase; catalytic subunit; MPST: mercaptopyruvate sulfurtransferase |
| GO:0000096 | Sulfur amino acid metabolic process | 6.35E-06 | 5.10E-03 | 8.9 | 3713 | 20 | 146 | 7 | CDO1: cysteine dioxygenase type 1; GCLM: glutamate-cysteine ligase; modifier subunit; GADL1: glutamate decarboxylase-like 1; MUT: methylmalonyl coa mutase; GCLC: glutamate-cysteine ligase; catalytic subunit; CBS: cystathionine-beta-synthase; MPST: mercaptopyruvate sulfurtransferase |
| GO:0070125 | Mitochondrial translational elongation | 2.50E-08 | 6.54E-05 | 5.28 | 3713 | 77 | 146 | 16 | MRPL22: mt ribosomal protein l22; MRPL2: mt ribosomal protein l2; MRPL9: mt ribosomal protein l9; TUFM: tu translation elongation factor; mt; MRPL10: mt ribosomal protein l10; MRPS2: mt ribosomal protein s2; MRPS26: mt ribosomal protein s26; MRPS9: mt ribosomal protein s9; MRPS6: mt ribosomal protein s6; DAP3: death associated protein 3; MRPL27: mt ribosomal protein l27; MRPS11: mt ribosomal protein s11; MRPL38: mt ribosomal protein l38; MRP63: mt ribosomal protein 63; MRPL21: mt ribosomal protein l21; MRPL3: mt ribosomal protein l3 |
| GO:0070126 | Mitochondrial translational termination | 1.00E-07 | 2.10E-04 | 5.16 | 3713 | 74 | 146 | 15 | MRPL22: mt ribosomal protein l22; MRPL2: mt ribosomal protein l2; MRPL9: mt ribosomal protein l9; MRPL10: mt ribosomal protein l10; MRPS2: mt ribosomal protein s2; MRPS26: mt ribosomal protein s26; MRPS9: mt ribosomal protein s9; MRPS6: mt ribosomal protein s6; DAP3: death associated protein 3; MRPL27: mt ribosomal protein l27; MRPS11: mt ribosomal protein s11; MRPL38: mt ribosomal protein l38; MRP63: mt ribosomal protein 63; MRPL21: mt ribosomal protein l21; MRPL3: mt ribosomal protein l3 |

| GO Term | Description | *P* | *q* | Enrichment | *N* | *B* | *n* | *b* | Genes |
|---|---|---|---|---|---|---|---|---|---|
| GO:0006415 | Translational termination | 2.50E-07 | 4.36E-04 | 4.83 | 3713 | 79 | 146 | 15 | MRPL22: mt ribosomal protein l22; MRPL2: mt ribosomal protein l2; MRPL9: mt ribosomal protein l9; MRPL10: mt ribosomal protein l10; MRPS2: mt ribosomal protein s2; MRPS26: mt ribosomal protein s26; MRPS9: mt ribosomal protein s9; MRPS6: mt ribosomal protein s6; DAP3: death associated protein 3; MRPL27: mt ribosomal protein l27; MRPS11: mt ribosomal protein s11; MRPL38: mt ribosomal protein l38; MRP63: mt ribosomal protein 63; MRPL21: mt ribosomal protein l21; MRPL3: mt ribosomal protein l3 |
| GO:0006414 | Translational elongation | 3.47E-07 | 5.18E-04 | 4.42 | 3713 | 92 | 146 | 16 | MRPL22: mt ribosomal protein l22; MRPL2: mt ribosomal protein l2; MRPL9: mt ribosomal protein l9; TUFM: tu translation elongation factor; mt; MRPL10: mt ribosomal protein l10; MRPS2: mt ribosomal protein s2; MRPS26: mt ribosomal protein s26; MRPS9: mt ribosomal protein s9; MRPS6: mt ribosomal protein s6; DAP3: death associated protein 3; MRPL27: mt ribosomal protein l27; MRPS11: mt ribosomal protein s11; MRPL38: mt ribosomal protein l38; MRP63: mt ribosomal protein 63; MRPL21: mt ribosomal protein l21; MRPL3: mt ribosomal protein l3 |
| GO:0006790 | Sulfur compound metabolic process | 7.01E-09 | 3.66E-05 | 4.31 | 3713 | 124 | 146 | 21 | PAPSS2: 3'-phosphoadenosine 5'-phosphosulfate synthase 2; GADL1: glutamate decarboxylase-like 1; GSR: glutathione reductase; B3GNT3: udp-glcnac:betagal beta-1;3-n-acetylglucosaminyltransferase 3; MUT: methylmalonyl coa mutase; SUCLG2: succinate-coa ligase; gdp-forming; beta subunit; GALNS: galactosamine (n-acetyl)-6-sulfate sulfatase; ETHE1: ethylmalonic encephalopathy 1; TSTD1: thiosulfate sulfurtransferase (rhodanese)-like domain containing 1; CDO1: cysteine dioxygenase type 1; SQRDL: sulfide quinone reductase-like (yeast); SLC25A1: solute carrier family 25 (mt carrier; citrate transporter); member 1; GCLM: glutamate-cysteine ligase; modifier subunit; CBS: cystathionine-beta-synthase; GCLC: glutamate-cysteine ligase; catalytic subunit; MPST: mercaptopyruvate sulfurtransferase; SLC25A10: solute carrier family 25 (mt carrier; dicarboxylate transporter); member 10; G6PD: glucose-6-phosphate dehydrogenase; GSTO1: glutathione s-transferase omega 1; MGST1: microsomal glutathione s-transferase 1; HSPA9: heat shock 70kda protein 9 (mortalin) |
| GO:0043624 | Cellular protein complex disassembly | 1.46E-06 | 1.91E-03 | 4.24 | 3713 | 90 | 146 | 15 | MRPL22: mt ribosomal protein l22; MRPL2: mt ribosomal protein l2; MRPL9: mt ribosomal protein l9; MRPL10: mt ribosomal protein l10; MRPS2: mt ribosomal protein s2; MRPS26: mt ribosomal protein s26; MRPS9: mt ribosomal protein s9; MRPS6: mt ribosomal protein s6; DAP3: death associated protein 3; MRPL27: mt ribosomal protein l27; MRPS11: mt ribosomal protein s11; MRPL38: mt ribosomal protein l38; MRP63: mt ribosomal protein 63; MRPL21: mt ribosomal protein l21; MRPL3: mt ribosomal protein l3 |
| GO:0022904 | Respiratory electron transport chain | 1.55E-05 | 1.01E-02 | 3.98 | 3713 | 83 | 146 | 13 | UQCRC2: ubiquinol-cytochrome c reductase core protein ii; UQCRFS1: ubiquinol-cytochrome c reductase; rieske iron-sulfur polypeptide 1; COX4I1: cytochrome c oxidase subunit iv isoform 1; NDUFA9: nadh dehydrogenase (ubiquinone) 1 alpha subcomplex; 9; 39kda; NDUFA10: nadh dehydrogenase (ubiquinone) 1 alpha subcomplex; 10; 42kda; NDUFC2-KCTD14: ndufc2-kctd14 readthrough; COX10: cytochrome c oxidase assembly homolog 10 (yeast); COX4I2: cytochrome c oxidase subunit iv isoform 2 (lung); AIFM2: apoptosis-inducing factor; mt-associated; 2; COX7B: cytochrome c oxidase subunit viib; NDUFV1: nadh dehydrogenase (ubiquinone) flavoprotein 1; 51kda; NDUFS2: nadh dehydrogenase (ubiquinone) fe-s protein 2; 49kda (nadh-coenzyme q reductase); ETFA: electron-transfer-flavoprotein; alpha polypeptide |

| GO Term | Description | *P* | *q* | Enrichment | *N* | *B* | *n* | *b* | Genes |
|---|---|---|---|---|---|---|---|---|---|
| GO:0032984 | Protein-containing complex disassembly | 1.47E-06 | 1.71E-03 | 3.6 | 3713 | 127 | 146 | 18 | MRPL22: mt ribosomal protein l22; MRPL2: mt ribosomal protein l2; MRPL9: mt ribosomal protein l9; MRPL10: mt ribosomal protein l10; MRPS2: mt ribosomal protein s2; MRPS26: mt ribosomal protein s26; CHMP2A: charged multivesicular body protein 2a; MTIF2: mt translational initiation factor 2; CHMP5: charged multivesicular body protein 5; MRPS9: mt ribosomal protein s9; MRPS6: mt ribosomal protein s6; DAP3: death associated protein 3; MRPS11: mt ribosomal protein s11; MRPL27: mt ribosomal protein l27; MRPL38: mt ribosomal protein l38; MRP63: mt ribosomal protein 63; MRPL21: mt ribosomal protein l21; MRPL3: mt ribosomal protein l3 |
| GO:0022900 | Electron transport chain | 3.27E-05 | 1.90E-02 | 3.32 | 3713 | 115 | 146 | 15 | UQCRC2: ubiquinol-cytochrome c reductase core protein ii; UQCRFS1: ubiquinol-cytochrome c reductase; rieske iron-sulfur polypeptide 1; GSR: glutathione reductase; COX4I1: cytochrome c oxidase subunit iv isoform 1; NDUFA9: nadh dehydrogenase (ubiquinone) 1 alpha subcomplex; 9; 39kda; NDUFA10: nadh dehydrogenase (ubiquinone) 1 alpha subcomplex; 10; 42kda; NDUFC2-KCTD14: ndufc2-kctd14 readthrough; RDH16: retinol dehydrogenase 16 (all-trans); COX10: cytochrome c oxidase assembly homolog 10 (yeast); COX4I2: cytochrome c oxidase subunit iv isoform 2 (lung); AIFM2: apoptosis-inducing factor; mt-associated; 2; NDUFV1: nadh dehydrogenase (ubiquinone) flavoprotein 1; 51kda; COX7B: cytochrome c oxidase subunit viib; NDUFS2: nadh dehydrogenase (ubiquinone) fe-s protein 2; 49kda (nadh-coenzyme q reductase); ETFA: electron-transfer-flavoprotein; alpha polypeptide |
| GO:0051186 | Cofactor metabolic process | 1.51E-06 | 1.57E-03 | 3.07 | 3713 | 182 | 146 | 22 | PGK1: phosphoglycerate kinase 1; HK1: hexokinase 1; TPI1: triosephosphate isomerase 1; GSR: glutathione reductase; MUT: methylmalonyl coa mutase; SUCLG2: succinate-coa ligase; gdp-forming; beta subunit; NDUFA9: nadh dehydrogenase (ubiquinone) 1 alpha subcomplex; 9; 39kda; ETHE1: ethylmalonic encephalopathy 1; COQ7: coenzyme q7 homolog; ubiquinone (yeast); SLC25A1: solute carrier family 25 (mt carrier; citrate transporter); member 1; ABCB6: atp-binding cassette; sub-family b (mdr/tap); member 6; COX10: cytochrome c oxidase assembly homolog 10 (yeast); GCLM: glutamate-cysteine ligase; modifier subunit; PRDX1: peroxiredoxin 1; BLVRB: biliverdin reductase b (flavin reductase (nadph)); MMADHC: methylmalonic aciduria (cobalamin deficiency) cbld type; with homocystinuria; TALDO1: transaldolase 1; GCLC: glutamate-cysteine ligase; catalytic subunit; G6PD: glucose-6-phosphate dehydrogenase; GSTO1: glutathione s-transferase omega 1; MGST1: microsomal glutathione s-transferase 1; HSPA9: heat shock 70kda protein 9 (mortalin) |

| GO Term | Description | *P* | *q* | Enrichment | *N* | *B* | *n* | *b* | Genes |
|---|---|---|---|---|---|---|---|---|---|
| GO:0055114 | Oxidation-reduction process | 2.20E-10 | 2.29E-06 | 2.98 | 3713 | 324 | 146 | 38 | UQCRC2: ubiquinol-cytochrome c reductase core protein ii; PGK1: phosphoglycerate kinase 1; ACAA2: acetyl-coa acyltransferase 2; TPI1: triosephosphate isomerase 1; UQCRFS1: ubiquinol-cytochrome c reductase; rieske iron-sulfur polypeptide 1; GSR: glutathione reductase; COX4I1: cytochrome c oxidase subunit iv isoform 1; TSTD1: thiosulfate sulfurtransferase (rhodanese)-like domain containing 1; CDO1: cysteine dioxygenase type 1; SQRDL: sulfide quinone reductase-like (yeast); COQ7: coenzyme q7 homolog; ubiquinone (yeast); COX10: cytochrome c oxidase assembly homolog 10 (yeast); SCP2: sterol carrier protein 2; COX4I2: cytochrome c oxidase subunit iv isoform 2 (lung); COX7B: cytochrome c oxidase subunit viib; GSTO1: glutathione s-transferase omega 1; TXN: thioredoxin; ETFA: electron-transfer-flavoprotein; alpha polypeptide; DHRS1: dehydrogenase/reductase (sdr family) member 1; HK1: hexokinase 1; HCCS: holocytochrome c synthase; NDUFA9: nadh dehydrogenase (ubiquinone) 1 alpha subcomplex; 9; 39kda; NDUFA10: nadh dehydrogenase (ubiquinone) 1 alpha subcomplex; 10; 42kda; HIGD1A: hig1 hypoxia inducible domain family; member 1a; NDUFC2-KCTD14: ndufc2-kctd14 readthrough; ETHE1: ethylmalonic encephalopathy 1; RDH16: retinol dehydrogenase 16 (all-trans); PRDX1: peroxiredoxin 1; BLVRB: biliverdin reductase b (flavin reductase (nadph)); RDH13: retinol dehydrogenase 13 (all-trans/9-cis); CBS: cystathionine-beta-synthase; AIFM2: apoptosis-inducing factor; mt-associated; 2; NDUFV1: nadh dehydrogenase (ubiquinone) flavoprotein 1; 51kda; SLC25A10: solute carrier family 25 (mt carrier; dicarboxylate transporter); member 10; NDUFS2: nadh dehydrogenase (ubiquinone) fe-s protein 2; 49kda (nadh-coenzyme q reductase); MGST1: microsomal glutathione s-transferase 1; G6PD: glucose-6-phosphate dehydrogenase; DHRS13: dehydrogenase/reductase (sdr family) member 13 |
| GO:0022411 | Cellular component disassembly | 2.88E-05 | 1.77E-02 | 2.83 | 3713 | 171 | 146 | 19 | MRPL22: mt ribosomal protein l22; MRPL2: mt ribosomal protein l2; MRPL9: mt ribosomal protein l9; MRPL10: mt ribosomal protein l10; MRPS2: mt ribosomal protein s2; CTSV: cathepsin v; MRPS26: mt ribosomal protein s26; CHMP2A: charged multivesicular body protein 2a; MTIF2: mt translational initiation factor 2; CHMP5: charged multivesicular body protein 5; MRPS9: mt ribosomal protein s9; MRPS6: mt ribosomal protein s6; DAP3: death associated protein 3; MRPS11: mt ribosomal protein s11; MRPL27: mt ribosomal protein l27; MRPL38: mt ribosomal protein l38; MRP63: mt ribosomal protein 63; MRPL21: mt ribosomal protein l21; MRPL3: mt ribosomal protein l3 |
| GO:0006091 | Generation of precursor metabolites and energy | 5.65E-05 | 3.10E-02 | 2.61 | 3713 | 195 | 146 | 20 | UQCRC2: ubiquinol-cytochrome c reductase core protein ii; GBAS: glioblastoma amplified sequence; PGK1: phosphoglycerate kinase 1; HK1: hexokinase 1; TPI1: triosephosphate isomerase 1; UQCRFS1: ubiquinol-cytochrome c reductase; rieske iron-sulfur polypeptide 1; GSR: glutathione reductase; COX4I1: cytochrome c oxidase subunit iv isoform 1; NDUFA9: nadh dehydrogenase (ubiquinone) 1 alpha subcomplex; 9; 39kda; NDUFA10: nadh dehydrogenase (ubiquinone) 1 alpha subcomplex; 10; 42kda; NDUFC2-KCTD14: ndufc2-kctd14 readthrough; RDH16: retinol dehydrogenase 16 (all-trans); COX10: cytochrome c oxidase assembly homolog 10 (yeast); ATP5F1: atp synthase; h+ transporting; mt fo complex; subunit b1; COX4I2: cytochrome c oxidase subunit iv isoform 2 (lung); AIFM2: apoptosis-inducing factor; mt-associated; 2; NDUFV1: nadh dehydrogenase (ubiquinone) flavoprotein 1; 51kda; COX7B: cytochrome c oxidase subunit viib; NDUFS2: nadh dehydrogenase (ubiquinone) fe-s protein 2; 49kda (nadh-coenzyme q reductase); ETFA: electron-transfer-flavoprotein; alpha polypeptide |

| GO Term | Description | *P* | *q* | Enrichment | *N* | *B* | *n* | *b* | Genes |
|---|---|---|---|---|---|---|---|---|---|
| GO:0044281 | Small molecule metabolic process | 1.48E-08 | 5.15E-05 | 2.28 | 3713 | 512 | 146 | 46 | UQCRC2: ubiquinol-cytochrome c reductase core protein ii; PGK1: phosphoglycerate kinase 1; PAPSS2: 3'-phosphoadenosine 5'-phosphosulfate synthase 2; ACAA2: acetyl-coa acyltransferase 2; GADL1: glutamate decarboxylase-like 1; TPI1: triosephosphate isomerase 1; LTA4H: leukotriene a4 hydrolase; GSR: glutathione reductase; SUCLG2: succinate-coa ligase; gdp-forming; beta subunit; MFN1: mitofusin 1; GALNS: galactosamine (n-acetyl)-6-sulfate sulfatase; CDO1: cysteine dioxygenase type 1; COQ7: coenzyme q7 homolog; ubiquinone (yeast); OSBPL1A: oxysterol binding protein-like 1a; PDK3: pyruvate dehydrogenase kinase; isozyme 3; GCLM: glutamate-cysteine ligase; modifier subunit; SCP2: sterol carrier protein 2; MMADHC: methylmalonic aciduria (cobalamin deficiency) cbld type; with homocystinuria; SLC16A1: solute carrier family 16 (monocarboxylate transporter); member 1; TALDO1: transaldolase 1; GCLC: glutamate-cysteine ligase; catalytic subunit; LDLRAP1: low density lipoprotein receptor adaptor protein 1; FARSA: phenylalanyl-trna synthetase; alpha subunit; GSTO1: glutathione s-transferase omega 1; TXN: thioredoxin; ETFA: electron-transfer-flavoprotein; alpha polypeptide; GBAS: glioblastoma amplified sequence; HK1: hexokinase 1; TMEM55B: transmembrane protein 55b; B3GNT3: udp-glcnac:betagal beta-1;3-n-acetylglucosaminyltransferase 3; MUT: methylmalonyl coa mutase; ATP6V1A: atpase; h+ transporting; lysosomal 70kda; v1 subunit a; NDUFA9: nadh dehydrogenase (ubiquinone) 1 alpha subcomplex; 9; 39kda; GIMAP7: gtpase; imap family member 7; QARS: glutaminyl-trna synthetase; SLC25A1: solute carrier family 25 (mt carrier; citrate transporter); member 1; RDH16: retinol dehydrogenase 16 (all-trans); EPRS: glutamyl-prolyl-trna synthetase; PFKFB4: 6-phosphofructo-2-kinase/fructose-2;6-biphosphatase 4; ATP5F1: atp synthase; h+ transporting; mt fo complex; subunit b1; FH: fumarate hydratase; RDH13: retinol dehydrogenase 13 (all-trans/9-cis); CBS: cystathionine-beta-synthase; MPST: mercaptopyruvate sulfurtransferase; SLC25A10: solute carrier family 25 (mt carrier; dicarboxylate transporter); member 10; G6PD: glucose-6-phosphate dehydrogenase |

**Dataset S1.** $F_{ST}$ values that quantify patterns of divergence between sulfidic and non-sulfidic evotypes. This table is a modified GFF annotation file from the *X. maculatus* reference genomes, including FST values for sliding windows (25 kb windows at 5 kb steps) across the entire nuclear genome. Window coordinate for each chromosome are indicated by the start (BIN_START) and end (BIN_END) position. In addition, of $F_{ST}$ (weighted and average), we also reported the number of SNPs (N_variants) per window.
*Due to its size, this dataset is provided as a separate Excel file (DatasetS1-FST_results.xlsx).*

**Dataset 2.** Results of local ancestry analyses. The table indicates what cactus (see Figure S7) each gene in the genome is assigned to.
*Due to its size, this dataset is provided as a separate Excel file (DatasetS2-Cacti_results.xlsx).*

**Dataset 3.** Results of gene expression analysis using EVE. The table includes the gene ID based on the *X. maculatus* reference genome annotation file as well as the gene name, accession ID, and protein annotations based on a BLAST search against the SwissProt database. The table also contains the results of branch tests that evaluated shifts in gene expression associated with the colonization of $H_2S$-rich habitats as implemented in EVE. We list the likelihood ratio test statistic ($LRT_\theta$), *P*-value, and false discovery rate (FDR). In addition, the table lists average gene expression individuals from sulfidic and nonsulfidic habitats (in FPKM) and an indication of the directionality of expression shifts upon colonization of $H_2S$-rich habitats (upregulation: +1; downregulation: -1). Note that genes with significant evidence for upregulation are highlighted in green, those with significant evidence for downregulation in red.
*Due to its size, this dataset is provided as a separate Excel file (DatasetS3-EVE_results.xlsx).*

**Dataset S4.** Results of analyses of molecular evolution using branch tests implemented in paml. The table includes the gene ID based on the *X. maculatus* reference genome annotation file as well as the gene name, accession ID, and protein annotations based on a BLAST search against the SwissProt database. The table also contains the results of branch tests contrasted rates of nonsynonymous to synonymous substitutions (dN/DS, also known as ω) between nonsulfidic lineages (background) and sulfidic lineages (foreground). We list P-value, false discovery rate (FDR), $\omega_{background}$, and $\omega_{foreground}$. Note that genes with significant evidence for positive selection are highlighted in yellow. Besides two subunits of COX, CYTB also exhibited evidence for elevated ω (see Figure S9 for more details). It is also noteworthy that ARNT2 is also under positive selection. This gene encodes a transcription factor involved in aryl hydrocarbon receptor (AhR) signaling, which is responsible for regulating responses to xenobiotics (73). For other genes with evidence for positive or relaxed selection, it remains unclear how they may be related to the evolution of $H_2S$ tolerance.
*Due to its size, this dataset is provided as a separate Excel file (DatasetS4-Molecular_evolution_results.xlsx).*

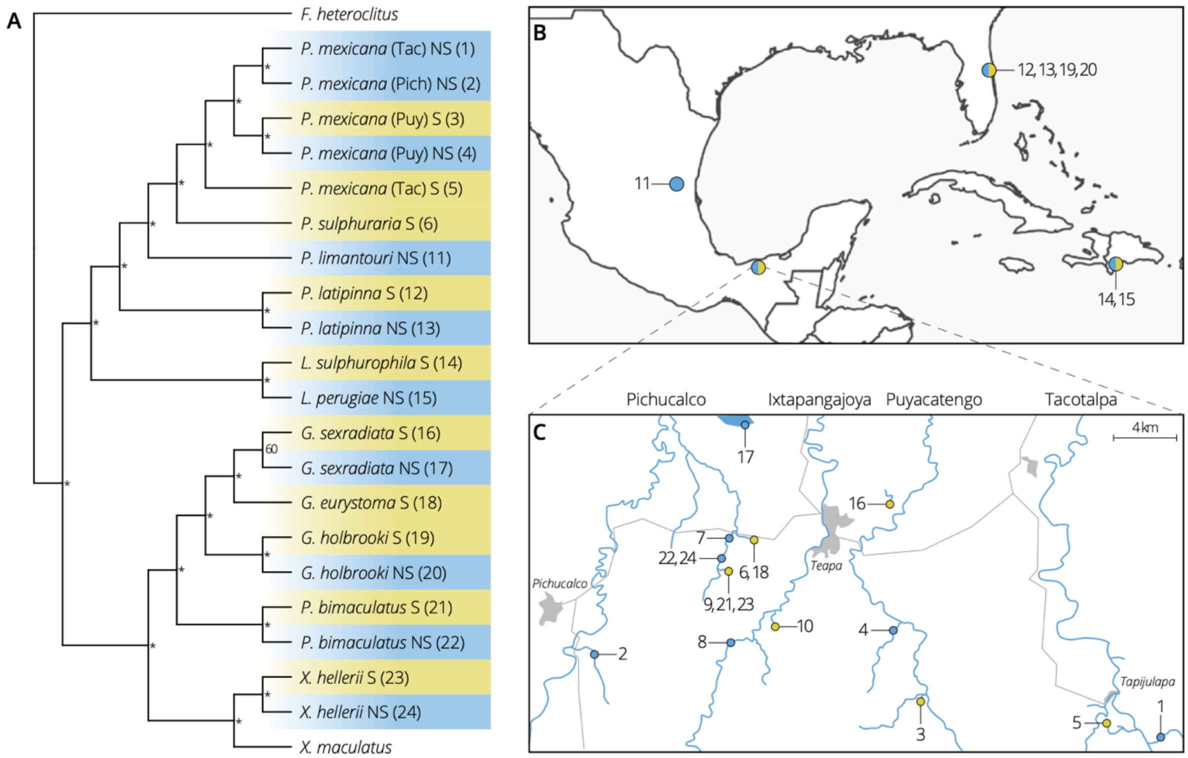


**Figure S1.** Overview of the phylogenetic relationships and distribution of lineages investigated in the comparative transcriptomics portion of this study. **A.** Phylogenetic tree of different sulfidic (S, underlined) and nonsulfidic lineages (NS), with *Fundulus heteroclitus* as outgroup. Asterisks indicate bootstrap support >0.99. **B.** Map depicting the collection locations of sulfide spring (yellow) and reference (blue) lineages. **C.** Detailed view of collection localities in southern Mexico; this map also includes sample sites from the comparative genomics analysis. Lineages included are numbered corresponding to Table S1. Black lines in panel C indicate major roads and gray shadings the location of major towns that have been added for orientation.

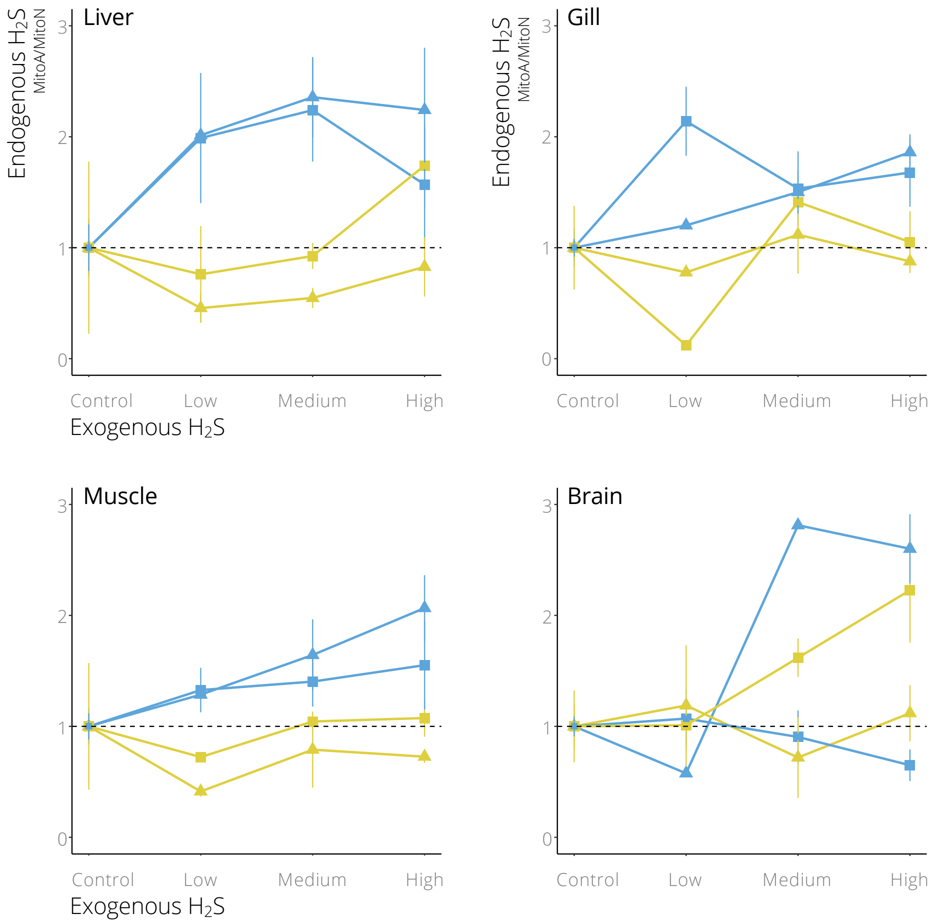


**Figure S2.** Relative change in endogenous $H_2S$ concentrations of live fish exposed to different levels of environmental $H_2S$, as quantified in different organs. For all graphs, yellow colors denote *P. mexicana* from sulfidic habitats, blue from nonsulfidic habitats. Symbols stand for populations from different river drainages (■: Tac; ▲: Puy). The results indicate that exposure to exogenous $H_2S$ leads to increases in endogenous $H_2S$ in fish from nonsulfidic environments. In contrast, fish from sulfidic environments have an increased ability to maintain low $H_2S$ endogenous concentrations upon exposure. These general trends are evident peripheral (gill) as well as internal organs (liver and muscle). However, these general patterns are much less clear in the brain, where there were no clear differences between sulfidic and nonsulfidic populations.

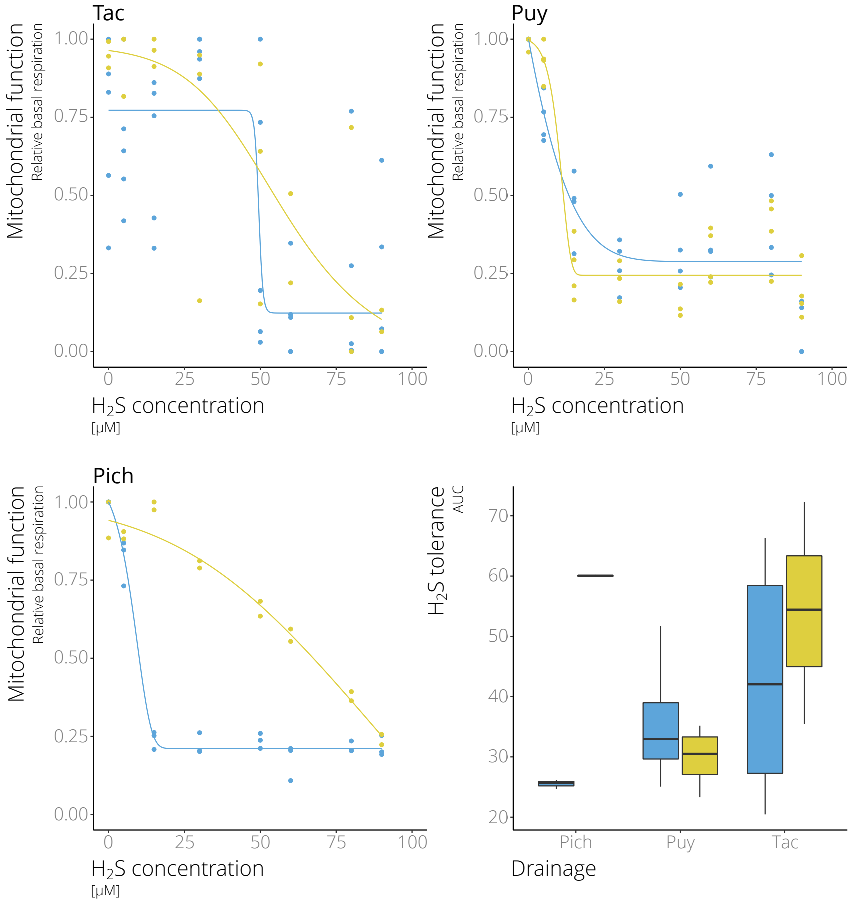


**Figure S3.** Dose response curves for basal mitochondrial respiration in sulfidic (yellow) and nonsulfidic (blue) populations of *P. mexicana* from different river drainages. Boxplots indicate levels of $H_2S$ tolerance (area under the curve, AUC) across populations. Variation in basal respiration was not related to any of the predictor variable (Table S8).

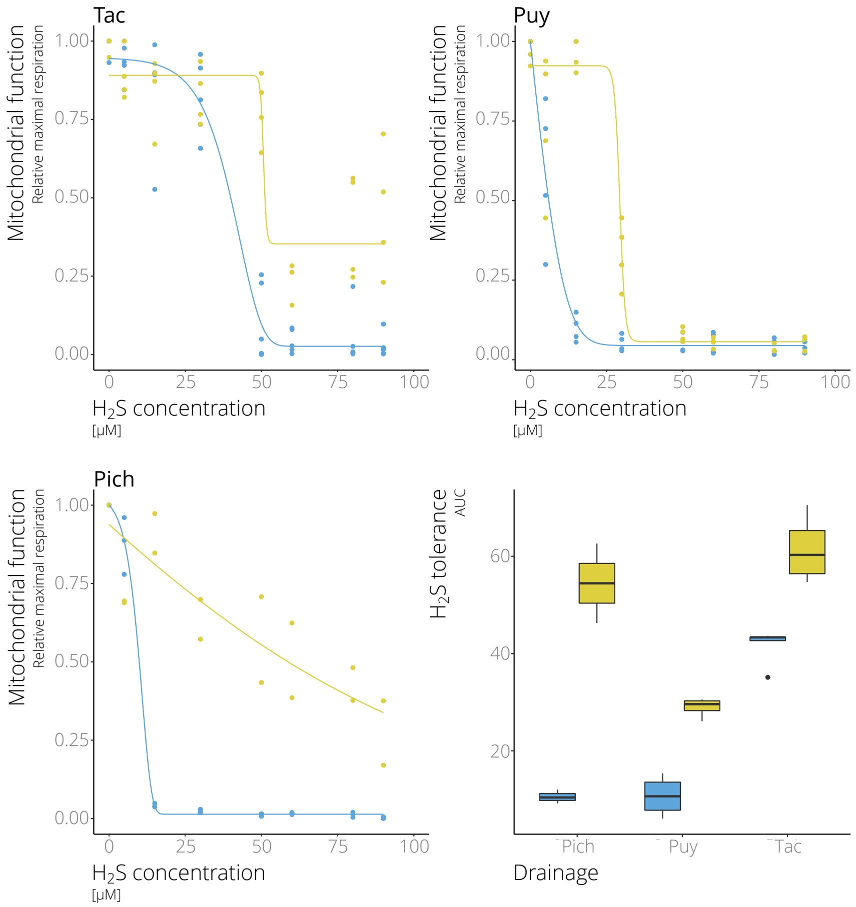


**Figure S4.** Dose response curves for maximal mitochondrial respiration in sulfidic (yellow) and nonsulfidic (blue) populations of *P. mexicana* from different river drainages. Boxplots indicate levels of $H_2S$ tolerance (area under the curve, AUC) across populations. The interaction between habitat type of origin and drainage of origin best explained variation in maximal respiration (Tables S9-S10).

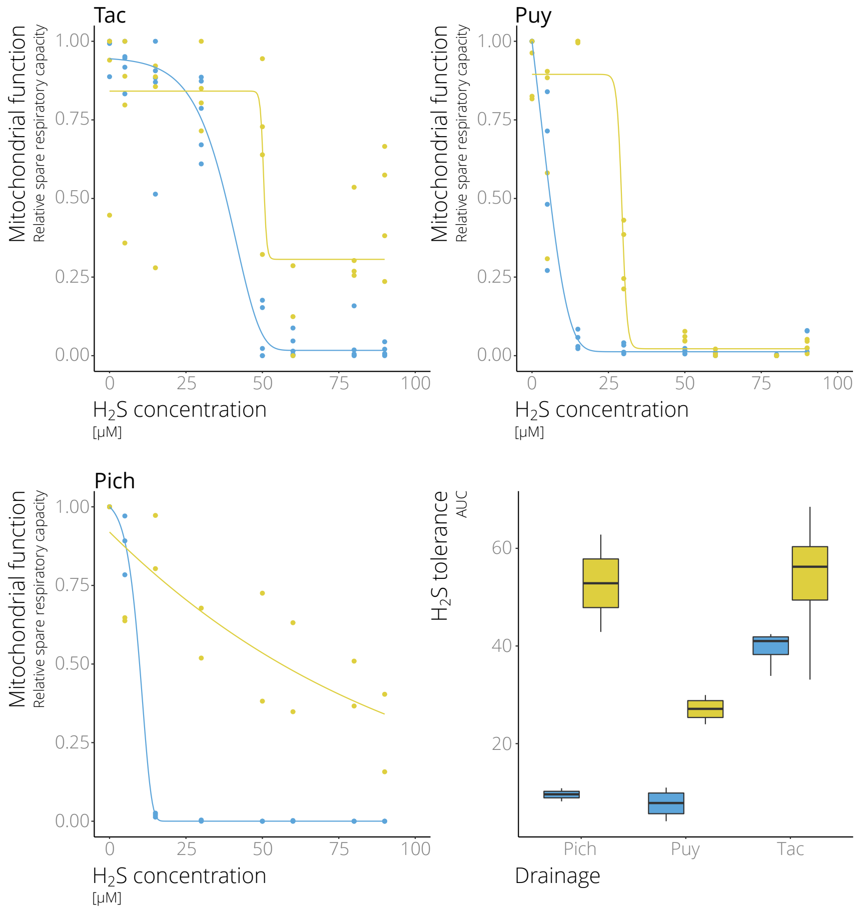


**Figure S5.** Dose response curves for spare respiratory capacity in sulfidic (yellow) and nonsulfidic populations (blue) of *P. mexicana* from different river drainages. Boxplots indicate levels of $H_2S$ tolerance (area under the curve, AUC) across populations. The interaction between habitat type of origin and drainage of origin best explained variation in spare respiratory capacity (Tables S11-S12).

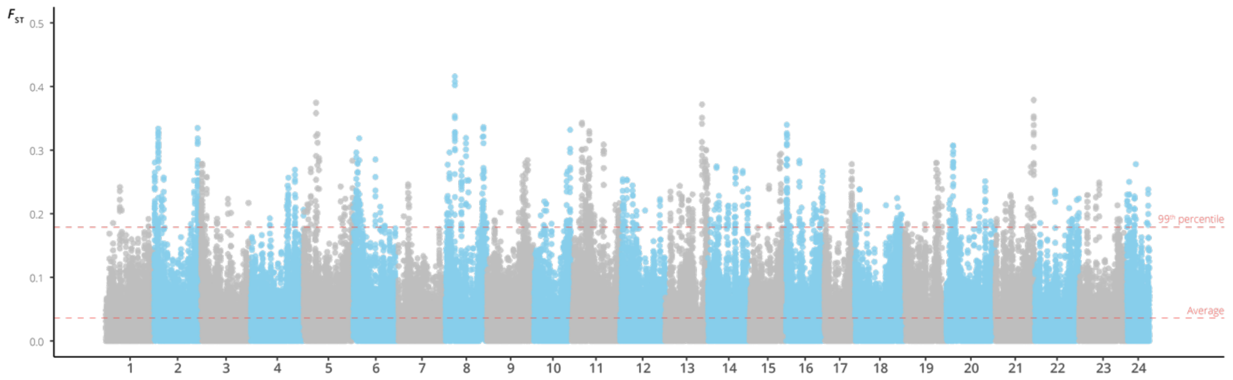


**Figure S6.** There is significant variation in local ancestry patterns across the genome of populations in the *P. mexicana* complex, as indicated by elevated $F_{ST}$-values between sulfidic and nonsulfidic populations (see Dataset S1 for details).

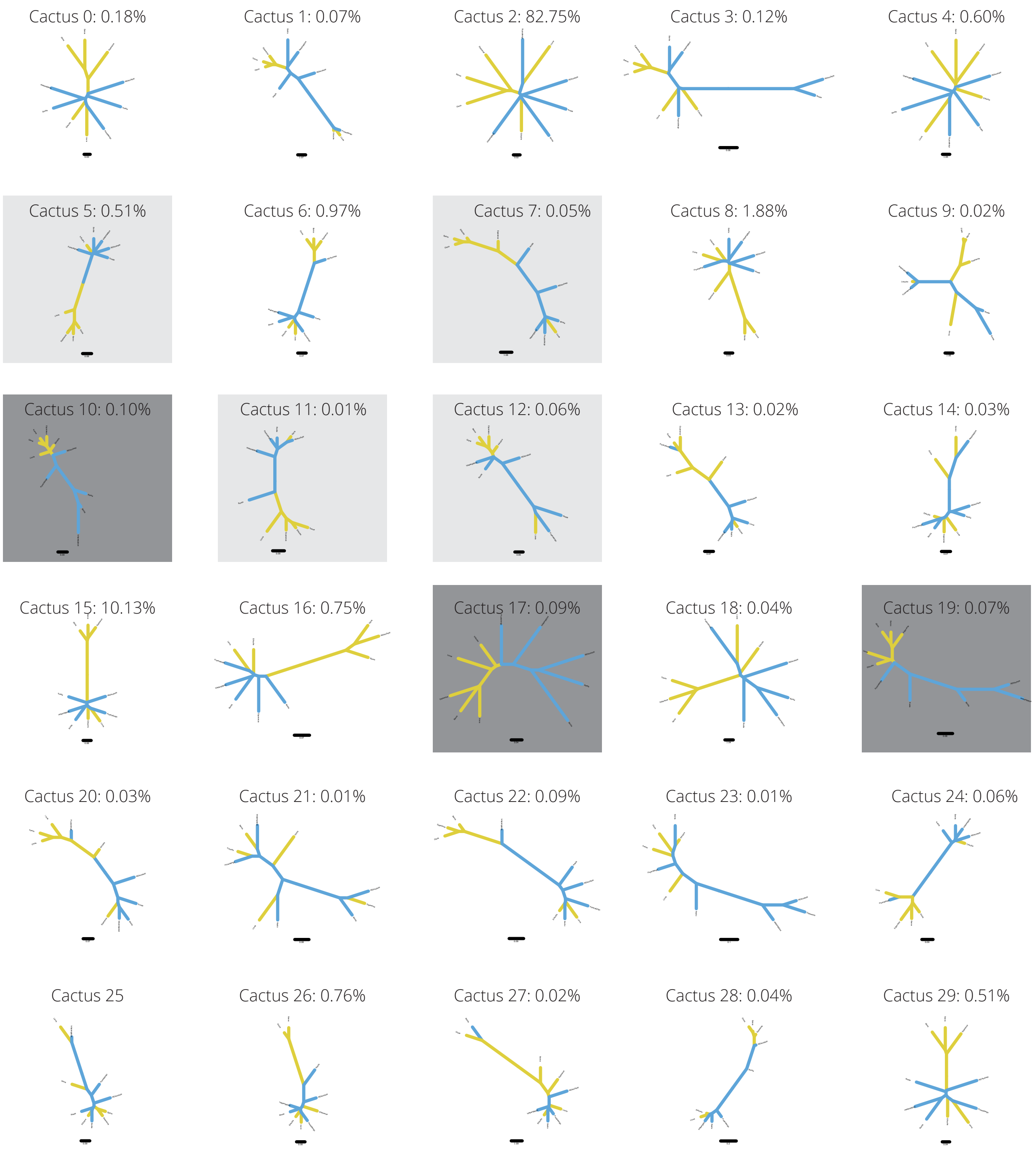


**Figure S7.** All 30 local topologies (cacti) hypothesized by Saguaro across the genome, including the percentage of the genome each cactus covered (lineages from sulfidic habitats are colored in yellow, those from non-sulfidic habitats in blue). Cacti 2 and 15 together covered over 90 % of the genome. Cacti 10, 17, and 19 (highlighted with dark gray boxes) exhibited strong clustering by ecotype, indicating monophyletic origin of putatively adaptive alleles. Cacti 5, 7, 11, and 12 (highlighted with light grey boxes) show clustering of four (out of five) sulfidic ecotypes. Assignment of genes to different cacti can be found in Dataset S2.

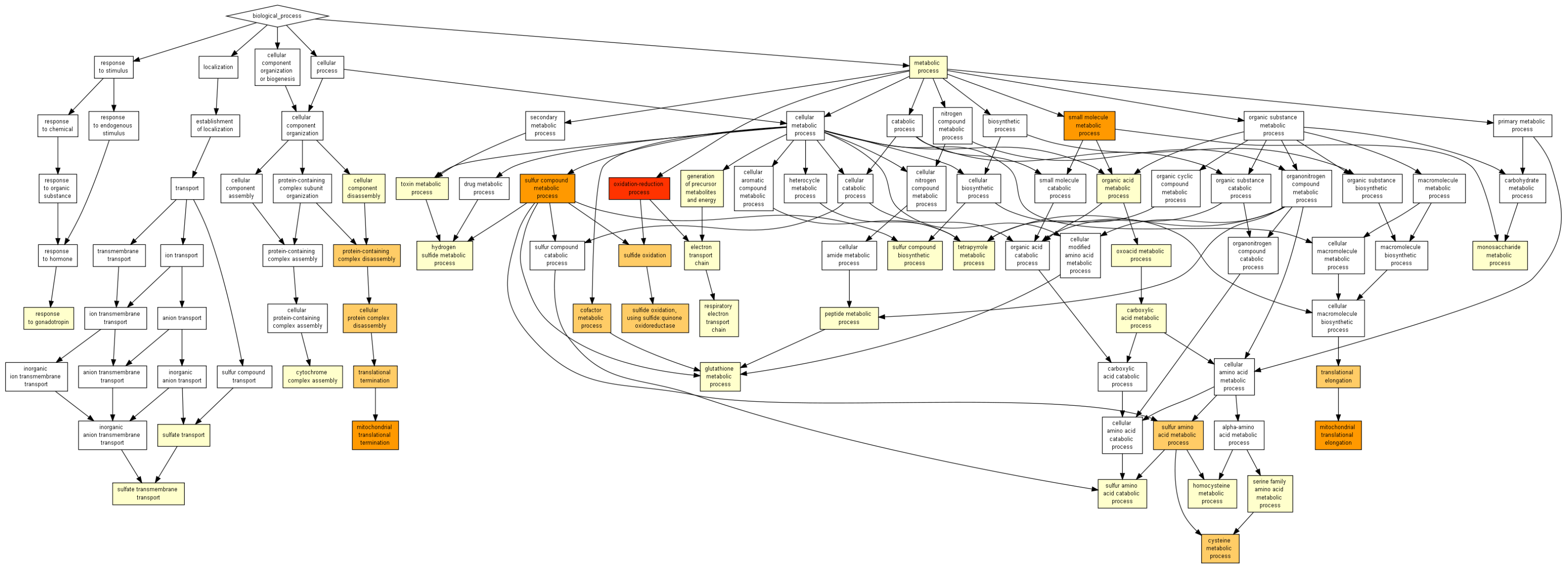


**Figure S8.** Overview of enriched GO terms (prior to FDR correction) in genes exhibiting convergent shift toward higher expression in lineages from $H_2S$-rich habitats (see Dataset S3 for details).

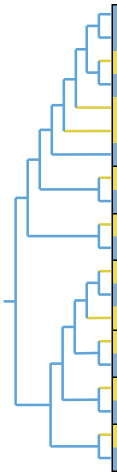

| CYTB | **13** | 37 | 42 | 43 | 57 | 61 | 89 | 109 | **118** | **170** | 185 | 214 | 215 | 240 | 243 | 304 | 306 | 313 | 315 | 333 | 347 | 348 | 353 | 356 | 360 |
|---|---|---|---|---|---|---|---|---|---|---|---|---|---|---|---|---|---|---|---|---|---|---|---|---|---|
| *Poecilia mexicana Tac NS* | | | I | | | M | | | | V | | | | I | A | | | | L | | | | I | | |
| *Poecilia mexicana Pich NS* | | | I | | | M | | | | V | | | | I | A | | | | L | | | | I | | |
| *Poecilia mexicanaPuy S* | | | I | | | M | | | | I | | | | I | A | | | | I | | | | I | | |
| *Poecilia mexicana Puy NS* | | | I | | | M | | | | V | | | | I | A | | | | L | | | | I | | |
| *Poecilia mexicana Tac S* | | | I | | | M | | | | V | | | | I | A | | | | L | | | | I | | |
| *Poecilia sulphuraria S* | | | V | | | S | | | | I | | | | V | T | | | | L | | | | M | | |
| *Poecilia limantouri NS* | | | I | | | M | | | | V | | | | I | A | | | | L | | | | I | | |
| *Poecilia latipinna S* | | | | | | | | | | | | | | | | | | | | | | | | | |
| *Poecilia latipinna NS* | | | | | | | | | | | | | | | | | | | | | | | | | |
| *Limia sulphurophila S* | | | | V | | | L | | | | | | | | | | | | | | | | | | |
| *Limia perugiae NS* | | | | A | | | M | | | | | | | | | | | | | | | | | | |
| *Gambusia sexradiata S* | I | | M | | | | | | I | V | L | | | | | V | F | | | | F | | | F | F |
| *Gambusia sexradiata NS* | V | | I | | | | | | I | V | F | | | | | I | F | | | | F | | | F | F |
| *Gambusia eurystoma S* | I | | M | | | | | | V | I | L | | | | | V | L | | | | Y | | | M | S |
| *Gambusia holbrooki S* | I | | | | | | | F | | | | | | | | | | | | | | V | | | |
| *Gambusia holbrooki NS* | V | | | | | | | Y | | | | | | | | | | | | | | I | | | |
| *Pseudoxiphophorus bimaculatus S* | | L | | | | | | | | | | D | A | | | | | | | L | | | | | |
| *Pseudoxiphophorus bimaculatus NS* | | F | | | | | | | | | | N | T | | | | | | | I | | | | | |
| *Xiphophorus hellerii S* | | | | | P | | | | V | | | | A | | | | | Q | | | | | | | |
| *Xiphophorus hellerii NS* | | | | | S | | | | I | | | | T | | | | | R | | | | | | | |

**Figure S9.** Amino acid differences in *CYTB* between lineages from sulfidic (yellow) and nonsulfidic (blue) habitats. Derived amino acids are shown in red. Bold letters indicate codons with convergent amino acid substitutions in different clades (separated by black horizontal lines) of sulfide spring fishes (codons 13, 118, and 170).
